# A General Approach to Adjusting Genetic Studies for Assortative Mating

**DOI:** 10.1101/2023.09.01.555983

**Authors:** Marta Bilghese, Regina Manansala, Dhruva Jaishankar, Jonathan Jala, Daniel J. Benjamin, Miles Kimball, Paul L. Auer, Michael A. Livermore, Patrick Turley

## Abstract

The effects of assortative mating (AM) on estimates from genetic studies has been receiving increasing attention in recent years. We extend existing AM theory to more general models of sorting and conclude that correct theory-based AM adjustments require knowledge of complicated, unknown historical sorting patterns. We propose a simple, general-purpose approach using polygenic indexes (PGIs). Our approach can estimate the fraction of genetic variance and genetic correlation that is driven by AM. Our approach is less effective when applied to Mendelian randomization (MR) studies for two reasons: AM can induce a form of selection bias in MR studies that remains after our adjustment; and, in the MR context, the adjustment is particularly sensitive to PGI estimation error. Using data from the UK Biobank, we find that AM inflates genetic correlation estimates between health traits and education by 14% on average. Our results suggest caution in interpreting genetic correlations or MR estimates for traits subject to AM.

## INTRODUCTION

Although it has been known for over 100 years that assortative mating (AM) can influence the results of genetic studies (Fisher, 1918), in recent years there has been increasing recognition of its potential impact in modern statistical genetic research (e.g., Border et al., 2022a, 2022b; Hartwig et al., 2018). Because results of genetic studies are simpler to interpret and less confounded when they are estimated in randomly mating populations, it would be useful to adjust estimates to what they would be under random mating. Theory-based adjustments for AM require strong assumptions about the sorting process or equilibrium state (Fisher, 1918; Crow and Feselstien, 1968; Crow and Kimura, 1970, Gianola 1982). As an alternative, we propose a general-purpose approach—requiring no assumptions about the sorting process or equilibrium—to evaluate how much of the genetic variance and genetic correlation is due to AM. Using the same principles, we also explore an adjustment to Mendelian randomization (MR) analyses that corrects for bias due to AM-induced LD arising from AM.

Our approach is based on three observations. First, regardless of the AM sorting process and history, AM affects estimates of genetic studies through its effects on linkage disequilibrium (LD). Second, because chromosomes segregate independently, the correlations between polygenic indexes (PGIs) constructed from single-nucleotide polymorphisms (SNPs) on different chromosomes identify the cross-chromosome component of AM-induced LD (Yengo et al., 2018). Third, combinatorial logic implies that the cross-chromosome component of AM-induced LD comprises the vast majority of all AM-induced LD. Therefore, using PGIs to estimate and adjust for cross-chromosome LD will purge nearly all bias due to AM.

Our approach can be applied to single traits to estimate how much the genetic variance is inflated by AM relative to the panmictic (i.e., random mating) genetic variance. Likewise, it can be applied to pairs of traits to assess how much the genetic correlation between traits is inflated relative to the panmictic genetic correlation. To illustrate, we study education and health-related traits, an important context with an expansive AM literature (e.g., Mare 1991, Domingue et al. 2014; Yengo et al. 2018). Applying our adjustment, we find that the genetic variance is inflated moderately for educational attainment (EA, by 10.1%, SE = 0.3%) and height (8.6%, SE = 0.3%) but little, if at all, for a range of health traits. The genetic correlation between EA and the health-related traits is inflated on average by less than 14%.

While our approach can be used to adjust genetic associations that are often used in MR studies, we find that it is inadequate for two reasons. First, in this setting, the adjustment is very sensitive to estimation error in the PGIs, producing additional biases to the MR estimates. Second, even if an error-free PGI were available, AM causes a selection bias in choosing which SNPs to use as instruments, which can generate upwardly biased MR estimates. We derive conditions under which we may expect to observe this bias. In simulations, we validate this theory and show that the latter bias, which we call *SNP-selection bias*, is often larger than the bias from AM-induced LD.

## RESULTS

### AM influences genetic studies through LD

We define AM very generally: AM is when mates have correlated traits or genotypes, regardless of how these correlations arise. For example, mates may sort directly on the trait of interest, or they may sort geographically across regions that have different mean levels of the trait (i.e., population stratification). For all types of AM, AM influences genetic studies through its effect on LD. Figure 1, Panel A illustrates AM-induced LD.

**Figure 1.**
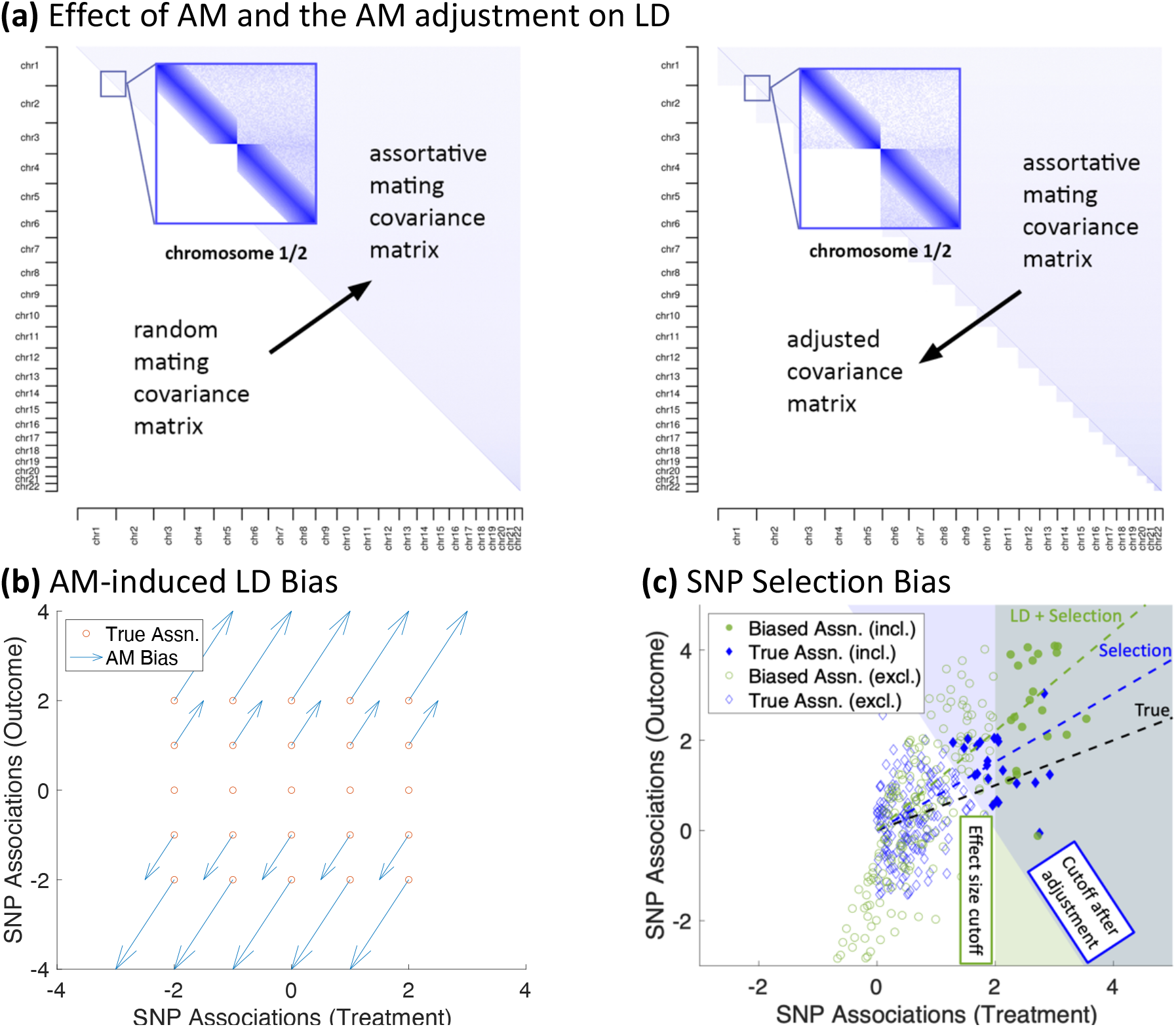
Conceptual illustration of assortative mating, the adjustment, and SNP selection bias. *Note*: **(a)** This figure is a stylized illustration of the effect of AM and our AM adjustment on LD in the genome. Each point in the plot represents the LD between a pair of SNPs. The darker shade of blue corresponds to stronger LD. The lower left portion of the left plot corresponds to a randomly mating population, with no LD except due to linkage. After many generations of AM, this transitions to the upper right, with low levels of genome-wide LD. The right figure illustrates the effect of our AM adjustment, with the upper-right portion representing AM as before, but the lower-left portion representing the LD structure implied by our adjustment, with no cross-chromosome LD, but with residual LD within each chromosome. **(b)** This figure illustrates how outcome AM biases GWAS associations for a pair of perfectly heritable traits when the genetic correlation between them is 0.5. The horizontal and vertical position of the red circles represent the true (i.e., panmictic) associations of a selected set of SNPs on the treatment and outcome traits, respectively. The blue arrows represent the how each of the SNP associations are affected due to direct sorting on the outcome (See Table 1 and Supplementary Materials). Figures corresponding to different sorting mechanisms and genetic correlations can be found in Supplementary Figures S2-S3. **(c)** This figure illustrates how SNP selection can bias MR studies. The green circles correspond to the SNPs, and the position of the circle represents its association with the treatment and outcome after AM. The solid circles (in the green shaded region) represent SNPs that have greater power in the treatment trait GWAS to be included in the MR study. The green dashed line is the line that passes through the origin and for which half of the solid green diamonds are above the line and half are below; the slope of this line is the (biased) MR-Median estimator. The blue diamonds correspond to true (panmictic) SNP associations. The solid diamonds in the blue shaded region correspond to the SNPs that would be included in an MR study if SNP selection was based on unadjusted estimates. The blue dashed line is the line that passes through the origin and for which half of the solid blue diamonds are above the line and half are below; the slope of this line is the MR-adjusted estimator which has corrected for LD bias but not SNP selection bias. The slope of the solid black line is the estimated effect of the treatment on the outcome if there were no SNP selection bias. In this figure, the SNPs have been oriented such that the true association on the treatment trait is positive.

Consider an additive model derived by projecting the trait onto the set of genotypes among individuals in a population. Formally, for a pair of traits, *Z* and *Y*:

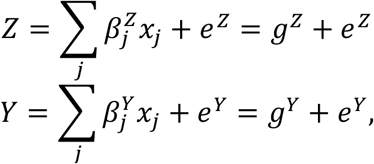

where *x*_*j*_ is the genotype of SNP *j* and *e*^*Z*^ and *e*^*Y*^ are the residuals of *Z* and *Y*, respectively. In this model, the SNP “effects” include not only causal effects but also confounds such as gene-environment correlation.

There are many ways that pairs of traits can be jointly analyzed in genetic studies; in this paper we consider genetic variance, genetic correlation, regressions with polygenic indexes (PGIs), and Mendelian randomization studies. The primary way that AM influences these analyses is by inducing LD between pairs of SNPs that would otherwise be uncorrelated in a randomly mating population. This LD arises because the maternally-inherited genotypes are correlated with the paternally-inherited genotypes, and these correlations can compound over many generations of AM. We consider the impact of AM-induced LD on each type of analysis below.

Genetic correlation can be defined in several ways (Okbay et al. 2016), the most common being the *effect correlation* (i.e., 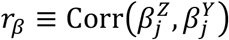) and the *score correlation* (i.e., 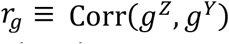) (Border et al. 2022). The effect correlation is unaffected by AM, but the score correlation can be. Because the focus of this paper is on AM, in what follows, we will use the term genetic correlation to mean score correlation.

The genetic correlation can be written:

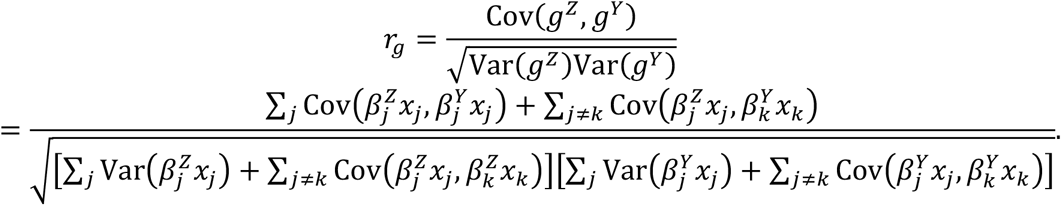

This expression contains many covariance terms corresponding to pairs of unlinked SNPs (e.g., SNPs on different chromosomes) that would be uncorrelated in a randomly mating population. However, in an AM population, these terms may be nonzero. Thus, AM can generate a genetic correlation between traits that would be uncorrelated in a randomly mating population.

AM influences regression analyses that include PGIs as covariates in a similar way as it affects genetic correlation. A PGI for trait *Z, ĝ*^*Z*^, may be thought of as a noisy, scaled estimate of the genetic factor for that trait: 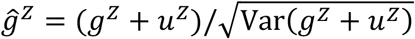with Cov(*g*^*Z*^, *u*^*Z*^) = Cov(*Y, u*^*Z*^) = 0, where *u*^*Z*^ represents the variation in the PGI due to estimation error of the PGI weights (Becker et al. 2021). So a regression of *Y* on *ĝ*^*Z*^ produces

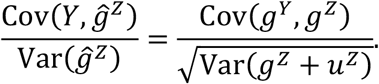

Because the genetic covariance is in the numerator of this expression, the same factors that influence genetic correlation will influence PGI regression coefficients.

There are two potential sources of bias due to AM in MR studies. The first is by inducing violations of MR’s assumptions. Most MR studies use GWAS summary statistics from a subset of SNPs to infer the causal effect of a “treatment” trait on an “outcome” trait. Formally, suppose the true causal model is

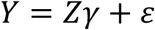

where *Y* is the outcome of interest (e.g., a health-related trait), *Z* is the treatment of interest (e.g., education), *γ* is the causal effect of the treatment of the outcome, and *ε* is the residual. Because Cov(*Z, ε*) may be nonzero, regression of *Y* on *Z* may produce a biased estimate of *γ*. However, MR tries to avoid this bias by taking the covariance of each side of this equation with respect to some SNP *x*_*j*_ and solving for *γ*. If the SNP genotype is uncorrelated with *ε*, then this produces

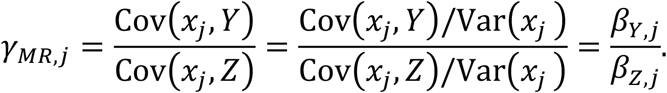

where *β*_*Y,j*_ and *β*_*Z,j*_ are the GWAS associations for SNP *j* for the outcome and treatment traits, respectively. In this paper, we refer to this ratio as the “MR ratio” for SNP *j*. The assumption that SNP genotype is uncorrelated with *ε* is called the “exclusion restriction.”

In most applications of MR, many SNPs are used as instruments, and there are a variety of MR methods that combine the MR ratios in different ways (see review in Sanderson et al., 2022). While inverse-variance-weighted MR requires that all SNPs included in the analysis satisfy the exclusion restriction, many of the MR methods have been developed to be robust to different ways that SNPs can violate the MR assumptions. For example, median-based MR (Bowden et al. 2016) is consistent if more than half of SNPs satisfy exclusion, and mode-based MR requires only that the modal SNP does (Guo et al., 2018). The MR-Egger method doesn’t require any SNP to satisfy exclusion, but it does require that the bias of the MR ratio is uncorrelated with the effect of the SNP on the treatment trait (Bowden et al. 2015). Researchers employing MR often apply a suite of approaches using the principle of “triangulation” (e.g., Pasman et al. 2018): that converging evidence across different MR methods with different assumptions provides stronger evidence of a causal effect.

Notice, however, that the exclusion restriction will only hold for SNP *j* if the SNP—and any genetic variant correlated with it—do not influence *ε*. This immediately makes apparent one way in which AM affects MR studies: AM can induce AM-induced LD between *every genetic variant*—not just SNPs—that influences the treatment. Therefore, in the presence of AM, *no* SNP will satisfy the exclusion restriction unless *all* genetic variants do. It follows that, in the presence of AM, the assumptions of median-based and mode-based MR are no weaker than those of inverse-variance weighted MR. Furthermore, because AM-induced LD between pairs of SNPs is stronger for SNPs that are more strongly associated with the treatment trait, MR-Egger estimates will also be biased.

A second way that AM can bias MR studies is by influencing the selection of SNPs to be included in the MR analysis. This mechanism is described in detail in the section *AM produces SNP selection bias in MR studies* below.

### Generalized AM theory has key implications

Descriptions of how AM-induced LD arises from AM and its expected influence on genetic studies has been explored previously in theory (Fisher, 1918; Crow and Feselstien, 1968; Crow and Kimura, 1970, Gianola 1982) and more recently in large-scale simulations (Hartwig et al. 2018, Border et al. 2022a, 2022b). While useful, the simulation evidence alone is limited in scope. Here, we derive theory that generalizes previous work (see Supplemental Note for details).

The sorting processes assumed in previous theory generally involved sorting on the particular trait or traits being studied. Our assumption subsumes these as special cases and builds on economic models of marital sorting (Becker 1973): females and males sort on latent traits, *S*, and *S*_-_, respectively, which may be a function of any number of observed or unobserved traits. Moreover, these functions may differ across males and females.

Now, consider projecting the sorting traits onto two observed traits, *Z* and *Y*, which (anticipating the MR application) we will respectively call the “treatment” and “outcome” traits:

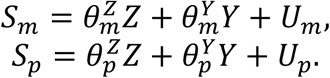

Then 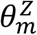 and 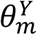 (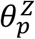 and 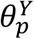) are the coefficients for predicting maternal (paternal) sorting by a linear combination of *Z* and *Y*, respectively, and *U*_*m*_ (*U*_*p*_) is the residual. The residual could include traits that are uncorrelated with *Z* and *Y*, nonlinearities, or components of the sorting traits that are not captured by *Z* and *Y*. We refer to 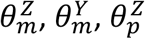, and 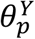 as the *sorting parameters* for the model. The magnitude of each sorting parameter can be thought of as representing how well the corresponding trait explains the sorting process for the corresponding sex.

We evaluated how genetic variances, genetic correlations, GWAS coefficients, and MR ratios evolve over time in this model (See Supplementary Note). The formulas for these expressions— which we validated in a series of simulations (see Online Methods, Supplementary Figures S4, S5, and S7, and Supplementary Tables S1-S2)—are complicated even when the sorting parameters are fixed over time and even after only one generation of sorting. However, the theory generates two simple conclusions that hold very generally. First, the expected effect of AM on these values is only a function of the sorting parameters, the panmictic heritabilities (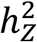 and 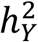) and genetic covariance/correlation (*ρ*_*ZY*_/*r*_*ZY*_) of *Z* and *Y*, and (in the case of GWAS coefficients) the association of the SNP with *Z, Y, U*_*m*_, and *U*_*p*_ (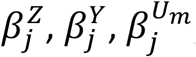, and 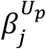, respectively). This result implies that the expected effect is *not* a function of other aspects of genetic architecture, such as the distribution of effect sizes or the fraction of SNPs that are truly pleiotropic. This is consistent with the conjecture of Border et al. (2022) based on simulation evidence. Second, the bias in GWAS coefficients is linear in 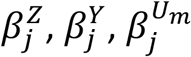, and 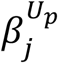. While the weights associated with each effect can be complex, the linearity is a useful property for understanding how biases can arise in MR.

In Table 1, we show how 3 simple forms of AM (AM on treatment, AM on outcomes, and cross-trait AM where mothers sort on the treatment and fathers on the outcome) affect several key parameters after a single generation. These special cases provide some intuition for the conditions under which we may or may not expect AM to affect our interpretation of genetic studies. For example, treatment AM will only affect the genetic variance of the outcome if the panmictic genetic correlation is nonzero. Also, all forms of AM will bias MR ratios except in knife-edge cases. This is inconsistent with the previous claims in the literature that pure single-trait AM on the treatment or outcome trait should not bias MR studies (e.g., Hartwig et al., 2018; Davies et al., 2019; Brumpton et al., 2020). These previous claims were based on simpler models and simulations than what we consider here. Panel B of Figure 1 also shows graphically what happens to SNP associations due to AM on the outcome trait when the treatment and outcome traits have a panmictic genetic correlation of 0.5. (Comparable figures for different settings are also found in Supplementary Figures S2-S3.)

**Table 1:** The effect of AM on certain genetic parameters in special cases.

|  | Excess<br>Variance (Z) | Excess<br>Variance (Y) | Genetic Correlation | GWAS<br>Bias (Z) | GWAS<br>Bias (Y) | MR Ratio |
| --- | --- | --- | --- | --- | --- | --- |
| Sorting only<br>on Z | $2\theta_m^Z \theta_p^Z h_Z^4$ | $2\theta_m^Z \theta_p^Z \rho_{ZY}^2$ | $r_{ZY} \sqrt{\frac{1 + 2\theta_m^Z \theta_p^Z h_Z^2}{1 + 2\theta_m^Z \theta_p^Z h_Z^2 r_{ZY}^2}}$ | $2\beta_j^Z \theta_m^Z \theta_p^Z h_Z^2$ | $2\beta_j^Z \theta_m^Z \theta_p^Z \rho_{ZY}$ | $\frac{\gamma + 2\rho_{ZY}}{1 + 2h_Z^2}$ |
| Sorting only<br>on Y | $2\theta_m^Y \theta_p^Y \rho_{ZY}^2$ | $2\theta_m^Y \theta_p^Y h_Y^4$ | $r_{ZY} \sqrt{\frac{1 + 2\theta_m^Y \theta_p^Y h_Y^2}{1 + 2\theta_m^Y \theta_p^Y h_Y^2 r_{ZY}^2}}$ | $2\beta_j^Y \theta_m^Y \theta_p^Y \rho_{ZY}$ | $2\beta_j^Y \theta_m^Y \theta_p^Y h_Y^2$ | $\gamma \frac{1 + 2h_Y^2}{1 + 2\gamma\rho_{ZY}}$ |
| Cross-trait<br>Sorting | $2\theta_m^Z \theta_p^Y h_Z^2 \rho_{ZY}$ | $2\theta_m^Z \theta_p^Y h_Y^2 \rho_{ZY}$ | $r_{ZY} \frac{1 + \theta_m^Z \theta_p^Y h_Z h_Y (r_{ZY} + 1/r_{ZY})}{1 + 2\theta_m^Z \theta_p^Y h_Z h_Y r_{ZY}}$ | $\beta_j^Z \theta_m^Z \theta_p^Y \rho_{ZY} + \beta_j^Y \theta_m^Z \theta_p^Y h_Z^2$ | $\beta_j^Z \theta_m^Z \theta_p^Y h_Y^2 + \beta_j^Y \theta_m^Z \theta_p^Y \rho_{ZY}$ | $\frac{\gamma + h_Y^2 + \gamma\rho_{ZY}}{1 + \gamma h_Z^2 + \rho_{ZY}}$ |

*Note:* Here we report how AM influences the excess genetic variance for Z and Y, the genetic correlation between Z and Y, the bias in a GWAS estimate of the effect of SNP $j$ on Z and Y, and the MR ratio after one generation of AM in three simple scenarios. The three sorting scenarios are (a) sorting only on Z ( $\theta_m^Y = \theta_p^Y = \beta_j^{U_m} = \beta_j^{U_p} = 0$ ), (b) sorting only on Y ( $\theta_m^Z = \theta_p^Z = \beta_j^{U_m} = \beta_j^{U_p} = 0$ ), and (c) cross-trait sorting ( $\theta_m^Y = \theta_p^Z = \beta_j^{U_m} = \beta_j^{U_p} = 0$ ). Excess variance is defined as the difference between the genetic variance the sorting population (after one generation) and the panmictic genetic variance. The GWAS bias is defined as the difference between the expected GWAS coefficient after just one generation of sorting and the expected GWAS coefficient in a randomly mating population. The MR Ratio is the expected MR Ratio in an infinite sample corresponding to a causal effect of $\gamma$ for a SNP that satisfies the exclusion restriction, such that $\beta_j^Y = \gamma\beta_j^Z$ . In the table above, $h_Z^2$ and $h_Y^2$ correspond to the panmictic heritabilities of Z and Y, respectively; and $\rho_{ZY}$ and $r_{ZY}$ correspond to the panmictic genetic covariance and genetic correlation, respectively, between Z and Y.

### A general AM adjustment genetic correlation

We propose a new method that estimates the fraction of the genetic correlation (and incidentally the fraction of the genetic variance) that is due to AM in order to better interpret the results of such studies. At a high level, our methods use the amount of cross-chromosome LD for PGIs as a proxy for the amount of AM-induced LD in the genome. While AM also influences LD within a chromosome, as illustrated in Figure 1, Panel A, the number of cross-chromosome SNP-pairs vastly outnumbers within-chromosome pairs. We have estimated that 94% of the AM-induced LD is captured by cross-chromosome LD for polygenic traits (See Online Methods). Therefore, if we can estimate the fraction of genetic correlation that is due to within-chromosome LD, that will be a good approximation of how much the genetic correlation is inflated by AM.

The fraction of the genetic correlation between a pair of traits that is due to within-chromosome LD is

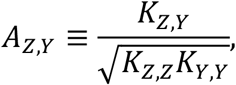

with

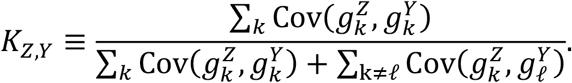

In these expressions, 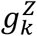 and 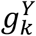 are the genetic factors of each trait for a chromosome *k*, which makes *K*_*Z,Y*_ the fraction of the covariance between a pair of traits that is due to within-chromosome LD. Likewise, *K*_*Z,Z*_ and *K*_*Y,Y*_ are the corresponding expressions for the variance. Thus *A*_*Z,Y*_ omits the portion of the genetic correlation due to cross-chromosome LD as a result of AM. Notice that under random mating, the cross-chromosome terms in the denominator are all expected to be zero, such that *K*_*Z,Y*_ = *K*_*Z,Z*_ = *K*_*Y,Y*_ = *A*_*Z,Y*_ = 1, as expected.

We approximate *K*_*Z,Y*_ using PGIs as

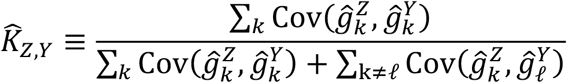

Where 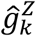 and 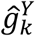 are the PGIs for each trait composed of only SNPs on chromosome *k*. Therefore, we obtain our estimate of *A*_*Z,Y*_ as

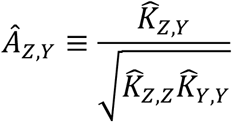

Multiplying an unbiased genetic correlation estimate by this value would adjust the estimate to what the genetic correlation would be in a randomly mating population. We obtain standard errors for this AM adjustment using a bootstrap procedure (see Online Methods).

The above adjustment would be perfect if 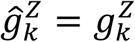 and 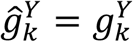. However, these equalities will not generally hold for two reasons: first, 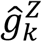 potentially omits some important genetic variants from 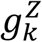 that may influence the trait; and second, 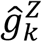 uses SNP weights that are estimated with error. We conducted a series of simulations with increasing amounts of error for a variety of genetic architectures and sorting mechanisms to evaluate the performance of our adjustment (see Online Methods). A summary of these results based on a randomly mating and a sorting population are found in Figure 2, Panels (a) and (b), respectively. In these simulations, *ϕ* represents the fraction of the variation in the PGI that is due to error.

**Figure 2:**
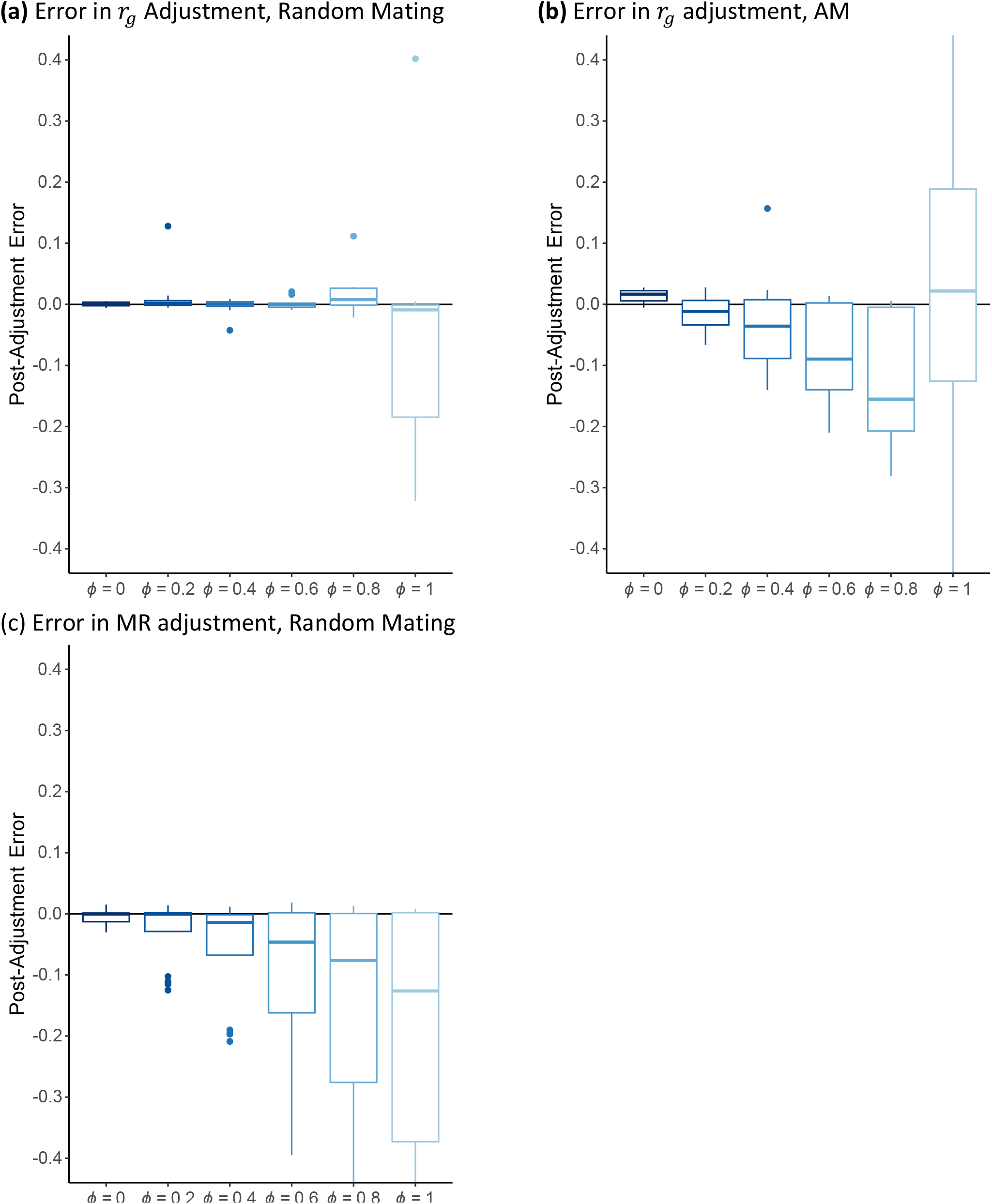
Summary of Error after AM Adjustment. *Note*: Box-plots of error in AM adjusted estimates across four genetic architectures (full pleiotropy/*r*_*g*_ = 0, full pleiotropy/ *r*_*g*_ = 0.5, partial pleiotropy, and no pleiotropy) simulated data for a randomly mating population. Each architecture was simulated 20 times. Full details on these simulations are found in the Online Methods. The parameter *ϕ* represents the fraction of the variation in the PGI that is due to error. **(a)** A box-plot for the error in AM-adjusted genetic-correlation estimates for different levels of PGI error under random mating. **(b)** A box-plot for the error in AM-adjusted genetic-correlation estimates for different levels of PGI error under AM. **(c)** A box-plot for the error in AM-adjusted genetic-correlation estimates for different levels of PGI error.

In the randomly mating population, the post-adjustment error appears mean zero for all levels of error, though the error variance becomes substantial for higher levels of *ϕ*. However, in the sorting population, there is a small amount of downward bias, representing an overcorrection. This implies that this adjustment will not identify a contribution of AM when there is none, but when noisy PGIs are used for the adjustment, if AM is truly driving some portion of a genetic correlation, our estimator produces an upper bound of the contribution of AM to genetic correlation. As a benchmark, *ϕ* = 0.4 the current PGI for educational attainment (see Online Methods). Complete results for this simulation can be found in Supplementary Figure S6 and Supplementary Table S3.

### An AM adjustment of GWAS associations

It is possible to adjust GWAS associations using a similar approach to the one described above. To see how, we use

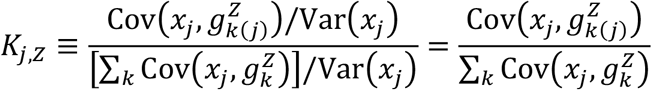

to denote the fraction of the GWAS associations for SNP *j* and trait *Z* that is due to cross-chromosome LD, where 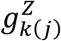is the genetic factor of *Z* for the chromosome that contains SNP *j*. As before, under random mating, the cross-chromosome covariance terms in the denominator are expected to be zero, and therefore *K*_*j,Z*_ = 1. As in the genetic correlation adjustment, we use PGIs to approximate this value:

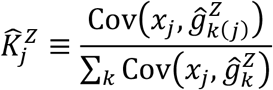

Using this approximation, the adjusted GWAS coefficients are

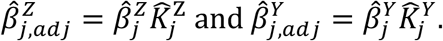

### MR adjustments are sensitive to PGI error

Adjusted GWAS coefficients can then be used to produce adjusted MR estimates. For example, adjusted MR-median estimates would be obtained by taking the ratio of the adjusted GWAS estimates and finding the median ratio. The adjusted MR estimate will be valid if the PGI is a good proxy for the true additive genetic factor. However, because the adjustment factor is a ratio of two small values, even small amounts of error in the PGI can lead to highly unstable adjusted GWAS estimates, and this instability is inherited by the MR estimate. In simulations summarized in Figure 2c (see Online Methods), we found that even a small amount of error leads to potentially large downward biases in the adjusted MR estimates even in a randomly mating population. (See Supplementary Figure S8 and Table S4 for complete results.)

### AM produces SNP-selection bias in MR studies

In addition to bias due to AM-induced LD inflating GWAS associations, AM can further lead to bias by how SNPs are selected for inclusion in MR analyses. Generally, in order to avoid weak instrument bias, only SNPs that meet some significance criteria are included in MR studies (e.g., each SNP must be associated with the “treatment” trait at *p* < 5 × 10^−8^). Under certain conditions, the SNPs that are best powered to meet this criterion are those that have large MR ratios. For example, consider a population that is sorting on just the *outcome* trait and where the outcome and treatment traits have a positive panmictic genetic correlation, *ρ*_*ZY*_ > 0. (This case is also illustrated in Panel (c) of Figure 1.) In this case, the bias of the GWAS association of SNP *j* for the *treatment* trait, 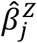, is

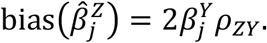

where 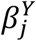 is the effect of SNP *j* on the outcome. Because the bias is largest when 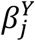 is largest, these SNPs will be more likely to reach the statistical threshold for inclusion. Consequently, more SNPs will be included with high values of 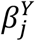 and therefore more SNPs with larger MR ratios, leading to additional bias away from zero in MR estimates.

This phenomenon can occur whenever the bias for the treatment GWAS estimate is a function of the effect of the SNP on the outcome trait, 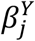. Under the sorting model described above, due to the linearity of the bias in GWAS, we show that this will be the case when

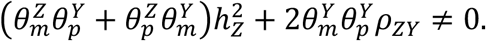

Intuitively, the first term will be non-zero if there is cross-trait AM and the treatment trait is heritable. The second term will be non-zero if there is single-trait AM on the outcome trait and the treatment and outcome traits are genetic correlated (as in the example described above).

### Real-data analysis

We used our adjustments to obtain estimates of how much AM affects the genetic variances and genetic correlations for education and health in real data. (Results related to MR analyses are included in Supplementary Table S7 but are omitted here due to their potential unreliability and instability.) The individual-level data for our real-data analysis come from the UK Biobank. Specifically, we used the *N* = 148,506 individuals from Tranche 1 of the SSGAC’s PGI Repository. All of these individuals have European genetic ancestries. The Educational Attainment PGI was based on Okbay et al. (2022). The health-related trait PGIs for our AM adjustment came from the SSGAC PGI Repository (Becker et al., 2021). The health traits we considered were asthma, BMI, cigarettes per day, ever smoker, depressive symptoms, drinks per week, neuroticism and self-rated health. We also included height as an additional health-related variable partly due to its association with childhood nutrition but also because it is a trait for which there is a large known degree of AM (Stuhlp et al., 2017) and for which a highly predictive PGI exists. The underlying samples used to generate these PGIs can be found in Becker et al. (2021). More information about these traits is in Supplementary Table S5.

### The effect of AM on EA and health

Figure 3 displays the estimated fraction of the genetic variances that is due to AM for EA and our set of health variables, as well as the fraction of the genetic correlations between education and health that are driven by AM. With a fraction of 10.1% (SE = 0.3%), the inflation of the EA genetic variance is larger than the inflation for any health variable. Indeed, with the exception of height (8.6%, SE = 0.3%) and self-rated health (5.1%, SE = 0.6%), the inflation of the genetic variance is less than 3% for all health traits.

**Figure 3:**
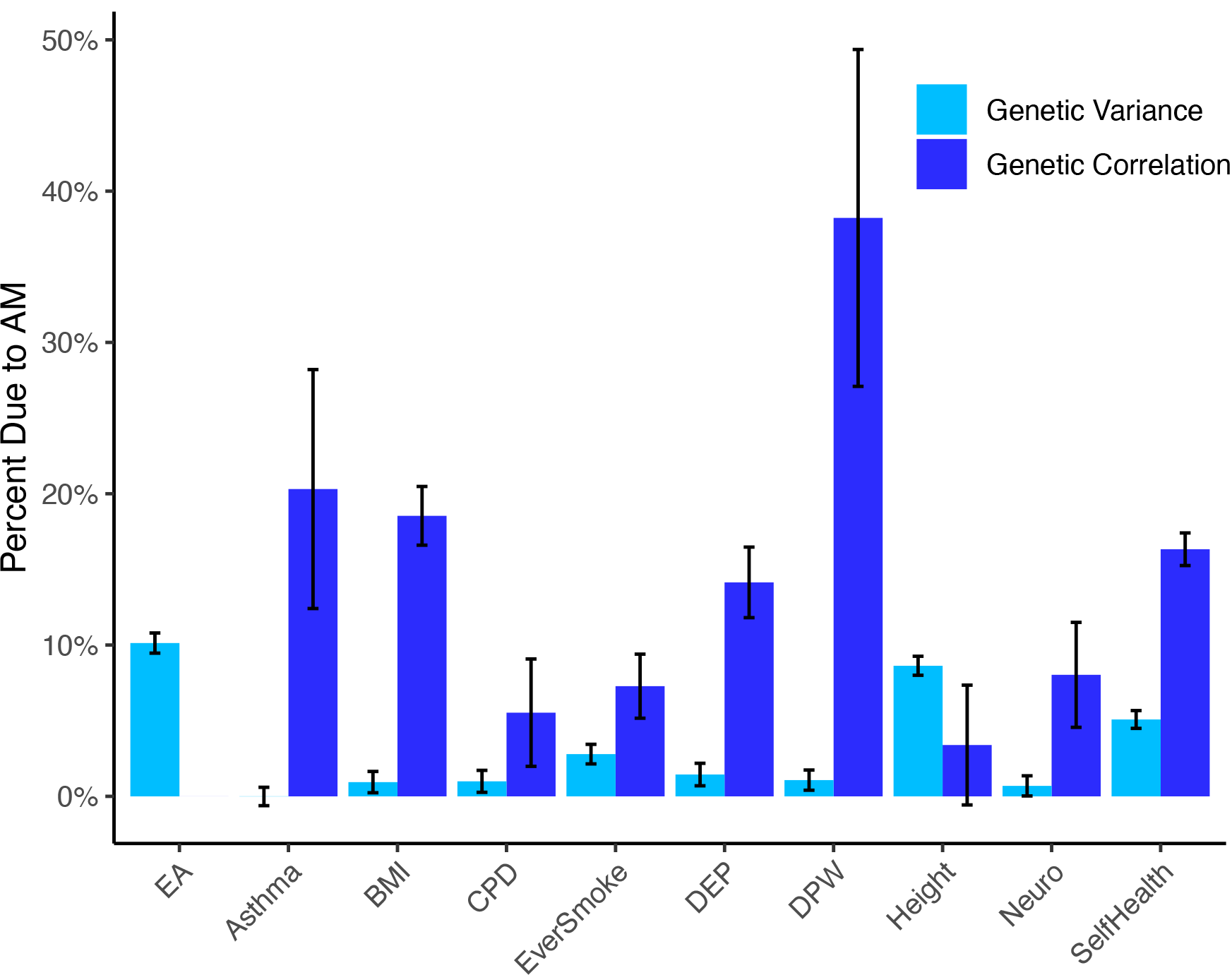
Fraction of Genetic Variance and Genetic Correlation with EA due to Assortative Mating. *Note*: Estimates of the fraction of genetic variance (light blue bars) and genetic correlation with EA (dark blue bars) that is due to AM. For the labeled health traits, the bar corresponds to the genetic correlation between educational attainment and the specified health traits. Data for this analysis comes from a subset of the UK Biobank (*N* = 148,506) and the PGIs come from the SSGAC’s PGI Repository (Becker et al. 2021). EA: Educational attainment; BMI: Body mass index; CPD: Cigarettes per day; DEP: Depressive symptoms; DPW: Drinks per week; Neuro: Neuroticism; Self Health: Self-rated health. Complete results can be found in Supplementary Table S6.

The inflation of the genetic correlations is more substantial. The largest inflation is seen for drinks-per-week, where 38% (SE = 11%) of the genetic correlation is estimated to be due to AM. The inflation of the EA-height genetic correlation is small and statistically weak (3%, SE = 4%), but for all other health traits, the inflation greater than 5%, with a mean across traits of 16%. However, due to noise in the PGIs used in the AM adjustment, these estimates should be considered an upper bound of the actual inflation. We further highlight that although some of the genetic correlation estimates cannot be explained by AM, this does not necessarily imply that there is a causal effect of EA on health (or vice versa) or even that there is a common underlying biological pathway since these results could also be explained by non-pleiotropic effects and local LD. However, for most of the health traits considered, we can rule out that the genetic correlation is primarily driven by AM.

## DISCUSSION

Much of modern genomics effectively ignores the role of AM, leading to potential misinterpretations of key results. Under random mating, these studies would have a much clearer interpretation. For this paper, we extended the existing theory on assortative mating and developed a general approach to adjust genetic studies to produce estimates equal to what they would have been in a randomly mating population.

The theoretical framework developed in this paper is based on behavioral sorting models from the economics literature (Becker 1973), which generalizes a broad class of sorting models commonly studied in genetics. A key contribution of our model is that mates can sort on a complex function of known and unknown traits, in contrast to a model where mates sort on a small number of known traits. As a generalization of existing work, our models are consistent with known theoretical results.

However, our theory also allowed us to fill in gaps missed by the extensive simulations presented previously, confirming some conjectures and refuting others. For example, based on a series of robustness analyses, Border et al. (2022) claim that the effect of AM on genetic correlation does not depend on the joint genetic architecture of the two traits (e.g., number of causal SNPs or whether causal SNPs are pleiotropic) or on the mate selection algorithm.

Consistent with this conjecture, we showed that the inflation of the genetic correlation is only a function of the sorting parameters, the panmictic heritabilities and genetic correlation, and no other higher order parameters. Our theory also refuted the common claim that pure single-trait AM on the treatment or outcome trait should not bias MR studies (e.g., Hartwig et al., 2018; Davies et al., 2019; Brumpton et al., 2020). The basis of this claim arises from a series of simulations that only evaluated cases where all SNPs that affected the treatment were valid instruments and with homogeneous treatment effects. Under weaker (and more plausible) assumptions, we showed that bias will arise even when there is only single trait sorting on either the treatment or outcome.

In addition to the consequences of AM described in detail above, the theory we derived may suggest other additional consequences of AM, potentially implicating any method that compares patterns of genetic effects across the genome. For example, some studies note the similarities in the associated biological pathways between pairs of traits (e.g., cognitive and noncognitive abilities (Demange et al., 2021) and psychiatric traits (Lynall et al. 2022)), concluding that the same types of cells mediate both traits. Such patterns could feasibly be driven by AM on one or both traits considered rather than a common biological pathway.

Relatedly, AM could drive the subtle genome-wide signals used to interpret genetic studies. Consider the Omnigenic Model (Boyle, Li, and Pritchard, 2017), which hypothesized that diffuse gene-regulatory mechanisms may explain a large fraction of heritability for all diseases. However, if these regulatory pathways are important to one or more highly polygenic traits on which people sort (such as height or EA), this could produce low levels of genome-wide LD. This could slightly inflate GWAS coefficients everywhere, including regulatory regions that are unrelated to the disease being studied, and could be misinterpreted as a weak omnigenic backbone.

That said, our empirical results on the genetic correlation between EA and health suggest that the situation may not be quite as dire as implied by Border et al. (2022). For the health traits they consider that overlap with our results (Neuroticism, Height, Ever smoker, Cigarettes per day, and BMI), they found that it is possible to produce genetic correlations that are on average 30% of the observed genetic correlations with only 5 generations of AM at observed levels. They concluded that “substantial variation in genetic correlation estimates can be explained by cross-mate phenotypic correlations.” In contrast we found that no more than 8.6% on average of the genetic correlation between EA and these traits is actually due to AM.

However, our theory showing potentially large AM-related biases underscores the growing recognition that more work needs to be replicated using family designs. Such studies break the AM-induced LD that arises from sorting, leaving only LD due to linkage. In such cases, however, caution is still warranted. For example, due to the lower power of family-based studies at currently available sample sizes, some MR studies select SNPs to use as instruments with population level data, while using family-based GWAS associations (which are free from AM-induced LD bias) in their MR analysis (Hwang et al., 2021). However, such an approach will still be subject to the SNP-selection problem we identify in this paper, leaving the results unreliable. In addition, while family data are able to correct for biases due to AM-induced LD, it cannot account for biases due to other violations of the exclusion restriction such as horizontal pleiotropy. Care should be taken to critically evaluate whether each SNP included is reliable (Burgess et al. 2019, Skrivankova 2021).

Future work may extend procedures like those proposed here, by accounting for within-chromosome LD and SNP selection bias, or by understanding how AM can also bias genetic analyses that we did not study. This work will be essential in efforts to better understand the complex interactions between genes, behavior, and health.

## ONLINE METHODS

### Between-chromosome AM-induced LD

AM can influence LD within a chromosome, but the contribution of within-chromosome SNP pairs to the total genetic (co)variance is expected to be negligible. Assuming that the *ex ante* expected contribution of any particular SNP pair to the genetic (co)variance is equal for all pairs, the expected contribution of just within-chromosome SNP pairs will be

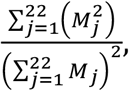

where *M*_*j*_ is the number of SNPs on chromosome *j*.Using the number of common SNPs (MAF > 0.01) in the European genetic ancestry subsample of the UK Biobank data, we calculated this number to be approximately 6%.

### Simulated data

We used a simple simulation framework to assess the effect of AM and the performance of our adjustment, which is summarized here. A more detailed description of the simulation procedure is found in the Supplementary Note. In our simulations, we began by simulating the genotypes of 2000 SNPs (spread across 20 equally sized chromosomes) for a population of 10,000 individuals. We assumed that the population had been randomly mating up until this generation such that each genotype for each person was drawn from a Binomial(2,*p*) distribution, where *p* = 0.5 is the allele frequency that was the same for every SNP.

We simulated two traits. We refer to as them as the “Treatment” and “Outcome” traits even when we conducted genetic correlation analyses and no causal relationship was assumed. We considered a variety of joint architectures with different levels of panmictic genetic correlation and different levels of pleiotropy. These are explained in greater detail in the Supplementary Note. To maximize the effect of AM and to maximize the statistical power of the analyses performed in the simulation, we simulated both traits to be perfectly heritable.

We considered 4 different mating mechanisms in our simulation: treatment AM, outcome AM, cross-trait AM, and multi-trait AM. Details about each of these mechanisms can be found in the Supplementary Note. To produce data for the subsequent generations, we first randomly split the data into males and females and matched them according to the mating mechanism. For example, for treatment AM, the male with the highest treatment trait value was matched with the corresponding female. Each pair produced two offspring. We assumed perfect recombination between SNPs such that the alleles inherited from each parent were independent. The traits of the children were constructed using the same effect sizes that were used for the parents. This procedure was repeated 3 times, using the children from one generation as the parents in the subsequent generation. Thus, we produced 4 generations of data: one corresponding to a randomly mating population, and 3 with increasing effects of sorting.

We used several sets of PGIs, each with a different amount of error. Specifically, the weight for SNP *j* and trait *Z* was set as

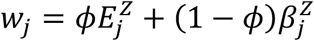

where *ϕ* ∈ {0.0, 0.2, 0.4, 0.6, 0.8, 1.0} is the parameter that sets the fraction of the PGI that is due to error, 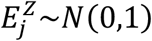 is an error term, and 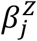 is the simulated effect size (also drawn from a standard normal distribution). The parameter *ϕ* is related to the parameter *ρ* from Becker et al. (2021) according to the following formula,

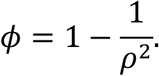

The interpretation of *ρ* is the amount of attenuation of the regression coefficient in a regression of a trait on its PGI relative to the coefficient that would be obtained if the additive SNP factor for the trait were used instead. Results for the performance of the adjustment when *ϕ* = 0 (the best-case scenario) are plotted in Supplementary Figures S5 and S7. Complete results showing the bias under all simulated scenarios are found in Supplementary Tables S3-S4.

In all simulations, we performed 20 replications of the simulation.

### Genetic correlation analysis in simulated data

Within each generation and for each genetic architecture and sorting mechanism, we calculated the genetic variance of each trait and the genetic correlation between them. In Supplementary Figure S4 and Supplementary Table S1, we report the ratio of the genetic variance in generations 2-4 relative to the genetic variance in generation 1. This is the amount of inflation of genetic variance due to AM. In Supplementary Figure S5 and Supplementary Table S1, we report the correlation of the genetic factors for the two traits in each generation.

We assessed the performance of the genetic correlation adjustment described in this paper by constructing sets of chromosome-specific “polygenic indexes” for each trait in each generation and calculating *A*_*Z,Y*_, which is the estimate of how much of the genetic correlation is due to within-chromosome LD. We then multiplied the genetic correlation in each generation by (1 – *A*_*Z,Y*_) to obtain an adjusted genetic correlation estimate.

### Mendelian randomization analysis in simulated data

To assess the effect of AM on MR analyses, we first performed a GWAS of the treatment and outcome traits in each generation. Because the simulated trait is quantitative, this was done by regressing the trait on each SNP (separately) and a constant using ordinary least squares. We selected the set of SNPs that had a p-value of less than 0.05/2000 = 2.5 × 10^7<^ to use in the MR analyses. We estimated three different MR estimators using the unadjusted GWAS results: MR-Egger, MR-Median, and MR-Mode. For the MR-Mode estimates, we used a rectangular kernel with a bandwidth of 0.1. More details on each of these methods are found in the Supplementary Materials.

We adjusted the GWAS summary statistics using the same PGIs described in the Online Methods section *Genetic correlation analysis in simulated data*, with various levels of error in the PGI. With the AM-adjusted GWAS results, we only applied the MR-Median estimator, which is more robust to the extreme draws sometimes observed when taking the ratio of two small values.

### Polygenic index weights

The polygenic indexes used in our real-data analysis of education and health came from the SSGAC’s Polygenic Index Repository (Becker et al. 2021). Specifically, we used the weights for Tranche 1 of the UK Biobank, which were built using a non-overlapping set of individuals from the residual UK Biobank and other cohorts. The weights were constructed using LDpred (Vilhjálmsson et al. 2015). See Becker et al. (2021) for more details on the split of the UK Biobank in three partitions and on the construction of the PGIs. We used these weights to make chromosome-specific PGIs using Plink2.0 (Chang et al. 2015).

### UK Biobank data

The genotypes used to make the PGIs for the real-data analysis came from the Tranche 1 of the UK Biobank (N=148,506).

The educational attainment variable we considered was the number of years of school completed, constructed as in Okbay et al. (2022). We considered nine health variables: asthma, BMI, cigarettes per day, ever smoker, depressive symptoms, drinks per week, height, neuroticism and self-rated health. Details on these trait definitions are found in Supplementary Table S5.

For the MR analyses, we performed a GWAS of each trait in the Tranche 1 sample of the UK Biobank (Becker et al. 2021). We kept only the 429 SNPs with F-statistics in the GWAS of education greater than 10. We will refer to this subset of SNPs as MR SNPs. As in the simulations, we used the MR-Egger, MR-Median, and MR-Mode estimators in this data. For MR-Egger, separately for each MR SNP, we regressed the health trait’s GWAS effect estimates on the educational attainment’s GWAS effect estimates and a constant using ordinary least squares. For MR-Median, for each MR SNP, we took the ratio of the health trait’s GWAS association and the education GWAS association (we refer to this ratio as the “MR Ratio”). The median of the MR ratios was used as the estimated effect of education on health. For MR-Mode, MR ratios were calculated, and a distribution was fit to the ratios using kernel regression with a rectangular kernel. We then adjusted the GWAS associations using PGIs for the same UKB subsample. After adjusting the summary statistics for AM, we applied the MR-Median method to get an AM-induced-LD-adjusted MR estimate of the effect of education on the health trait. We repeated this procedure for each health trait studied.

### Confidence intervals for real-data analyses

All confidence intervals in the real data analyses were constructed using a bootstrap approach.

For the genetic correlation analyses, we drew from the UKB sample (with replacement) a new sample of 148,506 individuals. With this bootstrap sample, we calculated the genetic correlation adjustment and saved the results. This was repeated 100 times. We report the 2.5^th^ and 97.5^th^ percentile of these replications as the 95% confidence interval.

For the MR analyses, we likewise drew a bootstrap sample of 148,506 individuals, and performed a GWAS in the bootstrap sample. We adjusted the GWAS estimates using PGIs from the bootstrap sample as well. Using the unadjusted and adjusted estimates, we applied our set of MR estimators as done in the MR analyses described in the previous section. This was repeated 100 times. We reported the 2.5^th^ and 97.5^th^ percentile of these replications as the 95% confidence interval.

## ACKNOWLEDGEMENTS

We thank Peter Visscher, Loic Yengo, Richard Border, Alex Young, Mike Goddard, Naomi Wray, and Robel Alemu for their helpful comments on this work. The research has also been conducted using the UKB Resource under application numbers 11425. Informed consent was obtained from UKB subjects. M.B., D.J.B., M.K., and P.T. were supported by Open Philanthropy (grant 010623-00001 to D.J.B.); M.B., M.K., and P.T by National Institute on Aging (NIA)/National Institutes of Health (NIH) (grant R21-AG067585 to P.T.); M.B., D.J.B, and P.T by the NIA/NIH (grants R24-AG065184 and R01-AG042568 to D.J.B.).

## DATA AVAILABILITY

Data from the UK Biobank, including the PGIs from the SSGAC’s PGI Repository, is available upon application to the UK Biobank.

## CODE AVAILABILITY

Upon publication, the scripts for carrying out the simulations and data analysis will be released in a public GitHub Repository.

## AUTHOR CONTRIBUTIONS

M.B., D.J.B, M.K., P.L.A., M.A.L, and P.T. designed and oversaw the study. M.B. was the study’s lead analyst, writing simulations and carrying out data analysis. R.M. and D.J. also contributed to data analysis. J.J. helped design and review the scripts written for this paper. All authors contributed to and critically reviewed the manuscript.

## COMPETING INTERESTS

The authors declare no competing interests.

## ADDITIONAL INFORMATION

Supplementary information is available for this paper. Correspondence and requests for materials should be addressed to.

## Notes

### Competing Interest Statement

The authors have declared no competing interest.

