## Supplementary Note for "A General Approach to Adjusting Genetic Studies for Assortative Mating"

### I. Extended Theory of Assortative Mating

In this section, we describe a model of assortative mating (AM) and how AM influences genome-wide association studies (GWAS) and (Mendelian randomization) MR estimates. First, we describe the model of sorting. Then we then calculate what the effect of AM is on the LD between genetic variants after one generation of sorting. We then extend that calculation to show how AM-induced LD influences the bias of GWAS estimates. This simple, single-generation derivation is useful to develop intuition for what factors are important for influencing LD and selection bias in MR studies. One conclusion of the derivation is that the bias in GWAS coefficients is a linear function of the (conditional) association of each SNP on the treatment, outcome, and other traits on which the population may be sorting. Finally, we show that, in a population that already has undergone an arbitrary amount of sorting, the bias from AM will always be a linear function of these associations.

*The model of sorting*

We consider a model of AM, where mothers and fathers mate based on a pair of latent phenotypes, where $S_m$ is the maternal phenotype and $S_p$ is the paternal phenotype. Specifically, we assume that parents will sort based on their rank-order of the latent phenotype. (That is, the female with the highest value of $S_m$ with mate the male with the highest value of $S_m$, and so forth.) This model of common, ordinal preferences follows the classic economic model of AM of Becker (1973). In contrast to existing models of sorting in the genetics literature where parents sort within or across single, observed traits (e.g., Gianola et al. 1982), this model is more general since $S_m$ and $S_p$ can be a function of any other variables and can also include a random component.

Because these latent phenotypes are ordinal (as opposed to cardinal phenotypes), the model will still hold under monotonic transformations of the phenotypes. Because of this, it is possible to transform $S_m$ and $S_p$ such that they are each normally distributed, each with a variance of one, and such that $S \equiv S_m = S_p$ whenever they correspond to matched parents. (The mean of the latent phenotypes will be fixed in a later step.)

Next, we consider how the sorting phenotypes relate to a pair of observed traits, $Z$ and $Y$. (We assume that these traits have been standardized to mean zero and variance one.) In what follows, we will often refer to $Z$ as the "treatment trait" and $Y$ as the "outcome trait", in anticipation of an MR application of this theory, though we do not assume a causal relationship between them unless otherwise noted. Imagine we project each sorting phenotype onto $Z$ and $Y$, giving us

$$S_m = \theta_m^Z Z + \theta_m^Y Y + U_m \quad (1)$$
$$S_p = \theta_p^Z Z + \theta_p^Y Y + U_p. \quad (2)$$

In this model, we allow for some of the sorting to be described by the treatment and outcome traits, but we also allow for residual terms, $U_m$ and $U_p$, which could include other traits relevant for sorting or for the random component of sorting. We have also allowed for the $\theta$-parameters to vary for mothers and fathers, which allows for different traits to matter differently for men and women.

Imagine that we project the sorting phenotypes onto the space spanned by some set of SNPs:

$$S_m = \sum_\ell \alpha_\ell^m \left(x_{m,\ell}^t + x_{m,\ell}^u\right) + e_m \tag{3}$$
$$S_p = \sum_\ell \alpha_\ell^p \left(x_{p,\ell}^t + x_{p,\ell}^u\right) + e_p, \tag{4}$$

where $x_{m,\ell}^t$ and $x_{m,\ell}^u$ are the alleles that are transmitted and untransmitted, respectively, to the mother's offspring and $x_{p,\ell}^t$ and $x_{p,\ell}^u$ are the corresponding values for the father. In this case, $\alpha_\ell^m$ and $\alpha_\ell^p$ are not the causal effects of the SNPs but simply the (conditional) associations of each SNP. The values $e_m$ and $e_p$ are the maternal and paternal residuals that aren't explained by SNPs. The mean of the residuals is fixed by the sorting process. Specifically, because $S_m = S_p$, it must be the case that $\mathrm{E}(S_m) = \mathrm{E}(S_p)$. One way for this to hold is if

$$\mathrm{E}(e_m) = \mathrm{E}\left[\sum_\ell \alpha_\ell^p \left(x_{p,\ell}^t + x_{p,\ell}^u\right)\right] = 2\sum_\ell \alpha_\ell^p \pi_\ell \tag{5}$$

and

$$\mathrm{E}(e_p) = \mathrm{E}\left[\sum_\ell \alpha_\ell^m \left(x_{m,\ell}^t + x_{m,\ell}^u\right)\right] = 2\sum_\ell \alpha_\ell^m \pi_\ell, \tag{6}$$

where $\pi_\ell$ is the allele frequency of SNP $\ell$. This way

$$\mathrm{E}(S) = 2\sum_\ell \left(\alpha_\ell^m + \alpha_\ell^p\right)\pi_\ell. \tag{7}$$

It will be the same in mothers and fathers since we are only considering SNPs on the autosomes, which are inherited independently of the sex chromosomes.

We can also project $Z$, $Y$, $U_m$, and $U_p$ onto these SNPs, giving us

$$Z = \sum_\ell \beta_\ell^Z x_\ell + e^Z \tag{8}$$
$$Y = \sum_\ell \beta_\ell^Y x_\ell + e^Y \tag{9}$$
$$U_m = \sum_\ell \beta_\ell^{U_m} x_\ell + e^{U_m} \tag{10}$$
$$U_p = \sum_\ell \beta_\ell^{U_p} x_\ell + e^{U_p}. \tag{11}$$

Substituting these expressions into (1) and (2) and rearranging terms gives us

$$S_m = \sum_\ell \left(\theta_m^Z \beta_\ell^Z + \theta_m^Y \beta_\ell^Y + \beta_\ell^{U_m}\right) x_\ell + \theta_m^Z e^Z + \theta_m^Y e^Y + e^{U_m} \tag{12}$$
$$S_p = \sum_\ell \left(\theta_p^Z \beta_\ell^Z + \theta_p^Y \beta_\ell^Y + \beta_\ell^{U_p}\right) x_\ell + \theta_p^Z e^Z + \theta_p^Y e^Y + e^{U_p}. \tag{13}$$

Therefore, this implies that

$$\alpha_\ell^m = \theta_m^Z \beta_\ell^Z + \theta_m^Y \beta_\ell^Y + \beta_\ell^{U_m} \quad (14)$$
$$\alpha_\ell^p = \theta_p^Z \beta_\ell^Z + \theta_p^Y \beta_\ell^Y + \beta_\ell^{U_p}. \quad (15)$$

The intuition of these expressions is that the effect of some SNP on the sorting phenotypes is the sum of the effect of that SNP on the treatment, outcome, and other traits times the association of those traits with the sorting phenotype.

*AM-induced LD after one generation of AM*

We begin by assuming the set of parents come from a randomly mating population but that the parents themselves are not randomly mating. The key implication of this assumption is $\mathrm{Cov}(x_{m,j}^t, x_{m,k}^t) = 0$ for all $j$ and $k$. For the children however, we calculate

$$\begin{aligned}\mathrm{Cov}(x_j, x_k) &= \mathrm{Cov}(x_{m,j}^t + x_{p,j}^t, x_{m,k}^t + x_{p,k}^t) \\ &= \mathrm{Cov}(x_{m,j}^t, x_{m,k}^t) + \mathrm{Cov}(x_{m,j}^t, x_{p,k}^t) + \mathrm{Cov}(x_{p,j}^t, x_{m,k}^t) + \mathrm{Cov}(x_{p,j}^t, x_{p,k}^t) \\ &= \mathrm{Cov}(x_{m,j}^t, x_{p,k}^t) + \mathrm{Cov}(x_{p,j}^t, x_{m,k}^t).\end{aligned}$$

We now consider each of the remaining two terms in turn.

First, by the definition of covariance and conditional probabilities, we have

$$\begin{aligned}\mathrm{Cov}(x_{m,j}^t, x_{p,k}^t) &= \mathrm{E}(x_{m,j}^t x_{p,k}^t) - \mathrm{E}(x_{m,j}^t)\mathrm{E}(x_{p,k}^t) \\ &= \mathrm{E}(x_{m,j}^t x_{p,k}^t) - \pi_j \pi_k \\ &= \mathrm{P}(x_{m,j}^t = x_{p,k}^t = 1) - \pi_j \pi_k \\ &= \mathrm{P}(x_{m,j}^t = 1 | x_{p,k}^t = 1)\mathrm{P}(x_{p,k}^t = 1) - \pi_j \pi_k \\ &= \int \mathrm{P}(x_{m,j}^t = 1 | x_{p,k}^t = 1, S) f(S | x_{p,k}^t = 1) dS\, \pi_k - \pi_j \pi_k. \end{aligned} \quad (16)$$

Here, because mates are sorting on $S$, once we condition on $S$, $x_{p,k}^t$ contains no additional information about $x_{m,j}^t$. Therefore $\mathrm{P}(x_{m,j}^t = 1 | x_{p,k}^t = 1, S) = \mathrm{P}(x_{m,j}^t = 1 | S)$. Substituting this into (16) gives us

$$\begin{aligned}\mathrm{Cov}(x_{m,j}^t, x_{p,k}^t) &= \int \mathrm{P}(x_{m,j}^t = 1 | x_{p,k}^t = 1, S) f(S | x_{p,k}^t = 1) dS\, \pi_k - \pi_j \pi_k \\ &= \int \mathrm{P}(x_{m,j}^t = 1 | S) f(S | x_{p,k}^t = 1) dS\, \pi_k - \pi_j \pi_k \\ &= \int \frac{f(S | x_{m,j}^t = 1)\mathrm{P}(x_{m,j}^t = 1)}{f(S)} f(S | x_{p,k}^t = 1) dS\, \pi_k - \pi_j \pi_k \\ &= \int \frac{f(S | x_{m,j}^t = 1) f(S | x_{p,k}^t = 1)}{f(S)} dS\, \pi_j \pi_k - \pi_j \pi_k.\end{aligned}$$

By the central limit theorem and under the assumption that the association of each SNP on the sorting phenotypes is small, each of the random variables, $(S|x^t_{m,j}=1)$, $(S|x^t_{p,k}=1)$, and $(S)$, are approximately normally distributed with variance one. The mean of $(S|x^t_{m,j}=1)$ is

$$\begin{aligned}
\mathrm{E}(S|x^t_{m,j}=1) &= \mathrm{E}\left(\sum_{\ell}\alpha^m_\ell\left(x^t_{m,\ell}+x^u_{m,\ell}\right)+e_m|x^t_{m,j}=1\right)\\
&= \mathrm{E}\left(\alpha^m_j x^t_{m,j}+\sum_{\ell\neq j}\alpha^m_\ell x^t_{m,\ell}+\sum_{\ell}\alpha^m_\ell x^u_{m,\ell}+e_m|x^t_{m,j}=1\right)\\
&= \alpha^m_j+\sum_{\ell\neq j}\alpha^m_\ell\pi_\ell+\sum_{\ell}\alpha^m_\ell\pi_\ell+2\sum_{\ell}\alpha^p_\ell\pi_\ell\\
&= \alpha^m_j-\alpha^m_j\pi_j+\alpha^m_j\pi_j+\sum_{\ell\neq j}\alpha^m_\ell\pi_\ell+\sum_{\ell}\alpha^m_\ell\pi_\ell+2\sum_{\ell}\alpha^p_\ell\pi_\ell\\
&= \alpha^m_j\left(1-\pi_j\right)+2\sum_{\ell}\left(\alpha^m_\ell+\alpha^p_\ell\right)\pi_\ell\\
&= \alpha^m_j\left(1-\pi_j\right)+\mathrm{E}(S). \qquad (17)
\end{aligned}$$

Likewise,

$$\mathrm{E}(S|x^t_{p,k}=1)=\alpha^p_k(1-\pi_k)+\mathrm{E}(S). \qquad (18)$$

Substituting in the (approximate) distribution functions and simplifying gives us

$$\begin{aligned}
\mathrm{Cov}\left(x^t_{m,j},x^t_{p,k}\right) &\approx \int\sqrt{2\pi}e^{-\frac{1}{2}\left[\left(S-\alpha^m_j(1-\pi_j)-\mathrm{E}(S)\right)^2+\left(S-\alpha^p_k(1-\pi_k)-\mathrm{E}(S)\right)^2-\left(S-\mathrm{E}(S)\right)^2\right]}dS\,\pi_j\pi_k-\pi_j\pi_k\\
&= \int\sqrt{2\pi}e^{-\frac{1}{2}\left[\left(S-\alpha^m_j(1-\pi_j)-\alpha^p_k(1-\pi_k)-\mathrm{E}(y)\right)^2-2\left(\alpha^m_j\alpha^p_k(1-\pi_j)(1-\pi_k)\right)\right]}dS\,\pi_j\pi_k-\pi_j\pi_k\\
&= e^{\alpha^m_j\alpha^p_k(1-\pi_j)(1-\pi_k)}\int\sqrt{2\pi}e^{-\frac{1}{2}\left[\left(S-\alpha^m_j(1-\pi_j)-\alpha^p_k(1-\pi_k)-\mathrm{E}(y)\right)^2\right]}dS\,\pi_j\pi_k-\pi_j\pi_k\\
&= e^{\alpha^m_j\alpha^p_k(1-\pi_j)(1-\pi_k)}\pi_j\pi_k-\pi_j\pi_k\\
&\approx \left[1+\alpha^m_j\alpha^p_k\left(1-\pi_j\right)(1-\pi_k)\right]\pi_j\pi_k-\pi_j\pi_k\\
&= \alpha^m_j\alpha^p_k\pi_j\left(1-\pi_j\right)\pi_k(1-\pi_k).
\end{aligned}$$

(The fourth line follows since the integrand is a normal pdf, and the fifth line follows from the Taylor expansion of an exponential.)

Following the same steps, we can also show that

$$\mathrm{Cov}\left(x^t_{p,j},x^t_{m,k}\right)=\alpha^m_k\alpha^p_j\pi_j\left(1-\pi_j\right)\pi_k(1-\pi_k).$$

And therefore,

$$\begin{aligned}\text{Cov}(x_j, x_k) &= \text{Cov}(x^t_{m,j}, x^t_{p,k}) + \text{Cov}(x^t_{p,j}, x^t_{m,k}) \\ &= (\alpha^m_j \alpha^p_k + \alpha^m_k \alpha^p_j)\pi_j(1-\pi_j)\pi_k(1-\pi_k). \end{aligned} \tag{19}$$

From this expression, we see that the LD between a pair of SNPs increases as the association between either SNP and the sorting phenotype increases. Indeed, in the special case the mothers and fathers sort on the same underlying phenotype (i.e., $\alpha^m_\ell = \alpha^p_\ell \forall \ell$), the covariance is directly proportional to each SNP's association with the sorting phenotype.

*Bias in GWAS after one generation of AM*

Next, we are interested in how AM produces bias in GWAS estimates. We begin considering the bias for the treatment trait. Using $\hat{\beta}^Z_j$ to denote the GWAS estimate of the association of SNP $j$ on trait $Z$, we calculate

$$\begin{aligned}\text{Bias}^Z_j &= E(\hat{\beta}^Z_j - \beta^Z_j) \\ &= \frac{\text{Cov}(x_j, Z)}{\text{Var}(x_j)} - \beta^Z_j \\ &= \frac{\text{Cov}(x_j, \sum_k \beta^Z_k x_k + e^Z)}{\text{Var}(x_j)} - \beta^Z_j \\ &= \frac{\sum_k \beta^Z_k \text{Cov}(x_j, x_k)}{\text{Var}(x_j)} - \beta^Z_j \\ &= \frac{\sum_{k\neq j} \beta^Z_k \text{Cov}(x_j, x_k)}{\text{Var}(x_j)} \\ &= \frac{\sum_{k\neq j} \beta^Z_k (\alpha^m_j \alpha^p_k + \alpha^m_k \alpha^p_j)\pi_j(1-\pi_j)\pi_k(1-\pi_k)}{\text{Var}(x_j)} \\ &\approx \frac{\sum_{k\neq j} \beta^Z_k (\alpha^m_j \alpha^p_k + \alpha^m_k \alpha^p_j)\pi_j(1-\pi_j)\pi_k(1-\pi_k)}{\pi_j(1-\pi_j)} \\ &= \sum_{k\neq j} \beta^Z_k (\alpha^m_j \alpha^p_k + \alpha^m_k \alpha^p_j)\pi_k(1-\pi_k) \\ &= \alpha^m_j \sum_{k\neq j} \beta^Z_k \alpha^p_k \pi_k(1-\pi_k) + \alpha^p_j \sum_{k\neq j} \beta^Z_k \alpha^m_k \pi_k(1-\pi_k). \end{aligned}$$

(The approximation in the seventh line above is because $\text{Var}(x_j) > \pi_j(1-\pi_j)$ due to AM, but the inflation of the variance will be small relative to $\text{Var}(x_j)$ if $\alpha^m_j$ and $\alpha^p_j$ are small.) Next, substituting in eqs (14-15), we get

$$\begin{aligned}\text{Bias}^Z_j = &\left(\theta^Z_m \beta^Z_j + \theta^Y_m \beta^Y_j + \beta^{U_m}_j\right) \sum_{k\neq j} \beta^Z_k \left(\theta^Z_p \beta^Z_k + \theta^Y_p \beta^Y_k + \beta^{U_p}_k\right) \pi_k(1-\pi_k) \\ &+ \left(\theta^Z_p \beta^Z_j + \theta^Y_p \beta^Y_j + \beta^{U_p}_j\right) \sum_{k\neq j} \beta^Z_k (\theta^Z_m \beta^Z_k + \theta^Y_m \beta^Y_k + \beta^{U_m}_k)\pi_k(1-\pi_k)\end{aligned}$$

$$= \beta_j^Z \sum_{k \neq j} \left[ \left(\theta_m^Z \theta_p^Z + \theta_p^Z \theta_m^Z\right)\left(\beta_k^Z\right)^2 + \left(\theta_m^Z \theta_p^Y + \theta_p^Z \theta_m^Y\right)\beta_k^Z \beta_k^Y + \theta_m^Z \beta_k^Z \beta_k^{U_p} + \theta_p^Z \beta_k^Z \beta_k^{U_m} \right] \pi_k(1-\pi_k)$$
$$+\beta_j^Y \sum_{k \neq j} \left[ \left(\theta_m^Y \theta_p^Z + \theta_p^Y \theta_m^Z\right)\left(\beta_k^Z\right)^2 + \left(\theta_m^Y \theta_p^Y + \theta_p^Y \theta_m^Y\right)\beta_k^Z \beta_k^Y + \theta_m^Y \beta_k^Z \beta_k^{U_p} + \theta_p^Y \beta_k^Z \beta_k^{U_m} \right] \pi_k(1-\pi_k)$$
$$+\beta_j^{U_m} \sum_{k \neq j} \left( \theta_p^Z \left(\beta_k^Z\right)^2 + \theta_p^Y \beta_k^Z \beta_k^Y + \beta_k^Z \beta_k^{U_p} \right) \pi_k(1-\pi_k)$$
$$+ \beta_j^{U_p} \sum_{k \neq j} \left( \theta_m^Z \left(\beta_k^Z\right)^2 + \theta_m^Y \beta_k^Z \beta_k^Y + \beta_k^Z \beta_k^{U_m} \right) \pi_k(1-\pi_k).$$

We next use $h_Z^2 \equiv \mathrm{Var}(\sum_k \beta_k^Z x_k) \approx \sum_{k \neq j} \left(\beta_k^Z\right)^2 \pi_k(1-\pi_k)$ to denote the random-mating heritability of the treatment trait, $\rho_{ZY} \equiv \mathrm{Cov}(\sum_k \beta_k^Z x_k, \sum_k \beta_k^Y x_k) \approx \sum_{k \neq j} \beta_k^Z \beta_k^Y \pi_k(1-\pi_k)$ to denote the random-mating genetic covariance of the treatment and outcome traits, $\rho_{ZU_m} \equiv \mathrm{Cov}\left(\sum_k \beta_k^Z x_k, \sum_k \beta_k^{U_m} x_k\right) \approx \sum_{k \neq j} \beta_k^Z \beta_k^{U_m} \pi_k(1-\pi_k)$ to denote the random-mating genetic covariance of the treatment and maternal sorting trait (after residualizing for $Z$ and $Y$), and $\rho_{ZU_p} \equiv \mathrm{Cov}\left(\sum_k \beta_k^Z x_k, \sum_k \beta_k^{U_p} x_k\right) \approx \sum_{k \neq j} \beta_k^Z \beta_k^{U_p} \pi_k(1-\pi_k)$ to denote the random-mating genetic covariance of the treatment and paternal sorting trait (after residualizing for $Z$ and $Y$), all under random mating.

Recall, however, that $U_m$ and $U_p$ are the residuals from projecting $S_m$ and $S_p$, respectively, onto $Z$ and $Y$, which implies that $U_m$ and $U_p$ are uncorrelated with $Z$ and $Y$. While it's technically feasible for those variables to be (phenotypically) uncorrelated while being genetically correlated, genetic correlations nearly always have the same sign as phenotypic correlations (Antilla et al., 2018), so the genetic correlation is likely to be negligible. Therefore, we assume that $\rho_{ZU_m} = \rho_{ZU_p} = \rho_{YU_m} = \rho_{YU_p} = 0$. Substituting in these terms gives us.

$$\mathrm{Bias}_j^Z = \beta_j^Z\left[2\theta_m^Z \theta_p^Z h_Z^2 + \left(\theta_m^Z \theta_p^Y + \theta_p^Z \theta_m^Y\right)\rho_{ZY}\right] + \beta_j^Y\left[\left(\theta_m^Y \theta_p^Z + \theta_p^Y \theta_m^Z\right)h_Z^2 + 2\theta_m^Y \theta_p^Y \rho_{ZY}\right]$$
$$+\beta_j^{U_m}\left(\theta_p^Z h_Z^2 + \theta_p^Y \rho_{ZY}\right) + \beta_j^{U_p}\left(\theta_m^Z h_Z^2 + \theta_m^Y \rho_{ZY}\right). \tag{20}$$

By symmetry, the bias for the outcome trait is

$$\mathrm{Bias}_j^Y = \beta_j^Z\left[\left(\theta_m^Y \theta_p^Z + \theta_p^Y \theta_m^Z\right)h_Y^2 + 2\theta_m^Z \theta_p^Z \rho_{ZY}\right] + \beta_j^Y\left[2\theta_m^Y \theta_p^Y h_Y^2 + \left(\theta_m^Z \theta_p^Y + \theta_p^Z \theta_m^Y\right)\rho_{ZY}\right]$$
$$+\beta_j^{U_m}\left(\theta_p^Y h_Y^2 + \theta_p^Z \rho_{ZY}\right) + \beta_j^{U_p}\left(\theta_m^Y h_Y^2 + \theta_m^Z \rho_{ZY}\right). \tag{21}$$

Below, we note that we may see SNP-selection bias when the bias on the treatment GWAS estimates is a function of the effect of the SNP on the outcome trait ($\beta_j^Y$). Therefore, we should expect to see SNP-selection bias when

$$\left(\theta_m^Y \theta_p^Z + \theta_p^Y \theta_m^Z\right)h_Z^2 + 2\theta_m^Y \theta_p^Y \rho_{ZY} \neq 0. \tag{22}$$

From this expression, we see that there are two key ways that selection bias can arise. First, if there is cross- or multi-trait AM, then $\left(\theta_m^Y\theta_p^Z + \theta_p^Y\theta_m^Z\right) \neq 0$ (except in rare knife edge cases), which implies that selection will always be a risk in that case. Second, even if there is no sorting on the treatment, there can be selection bias if there is sorting on the outcome and the treatment trait is genetically correlated with either the outcome trait or the residual of the selection trait. Notably, if there is no selection on the outcome trait (i.e., $\theta_m^Y = \theta_p^Y = 0$), then there is no selection bias.

Hartwig et al. (2018) claims that MR is not biased if there is single trait sorting solely on the treatment or outcome trait and if the traits are genetically uncorrelated. Eqs. (20) and (21) show that this is not true. Sorting on just the treatment implies that $\theta_m^Z = \theta_p^Z = 1$ and $\theta_m^Y = \theta_p^Y = \beta_j^{U_m} = \beta_j^{U_p} = 0$. Assume that the exclusion restriction is satisfied for SNP $j$ in a randomly mating population, such that $\beta_j^Y = \gamma\beta_j^Z$ where $\gamma$ is the causal effect of the treatment on the outcome. The MR ratio for this SNP (under the assumption of sorting only on treatment and no genetic correlation) is

$$\text{MRratio} = \frac{\beta_j^Y}{\beta_j^Z + 2\beta_j^Z h_Z^2} = \frac{\gamma\beta_j^Z}{\beta_j^Z + 2\beta_j^Z h_Z^2} = \frac{\gamma}{1+2h_Z^2}. \tag{23}$$

This will not equal $\gamma$ unless the treatment is not heritable (i.e., $h_Z^2 = 0$) or the treatment has not causal effect on the outcome (i.e., $\gamma = 0$). So the MR estimate will generally be biased even if the MR study only includes SNPs that satisfy exclusion under random mating.

Consider as well the case where parents only sort on the outcome trait. Then $\theta_m^Y = \theta_p^Y = 1$ and $\theta_m^Z = \theta_p^Z = \beta_j^{U_m} = \beta_j^{U_p} = 0$. The MR ratio for SNP $j$ in this case is

$$\text{MRratio} = \frac{\beta_j^Y + 2\beta_j^Y h_Y^2}{\beta_j^Z} = \frac{\gamma\beta_j^Z + 2\gamma\beta_j^Z h_Y^2}{\beta_j^Z} = \gamma\left(1 + 2h_Y^2\right). \tag{24}$$

This is also not generally equal to $\gamma$ unless the outcome is not heritable or the treatment has no causal effect on the outcome.

*Inflation of genetic variance and genetic correlation after one generation of AM*

While extensive theory has been derived for how AM affects genetic variance and genetic correlation (Fisher, 1918; Crow and Feselstien, 1968; Crow and Kimura, 1970, Gianola 1982), here we provide a parallel derivation in context of the parameterization above.

We define the "excess variance" as the difference in the genetic variance in the current generation and the genetic variance under random mating. That is, for the treatment trait,

$$\text{ExcessVariance}^Z = \text{Var}\left(\sum_j x_j\beta_j^Z\right) - \sum_j \text{Var}\left(x_j\beta_j^Z\right). \tag{25}$$

We further calculate

$$\text{ExcessVariance}^Z = \sum_j \sum_{k\neq j} \beta_j^Z \beta_k^Z \text{Cov}(x_j, x_k)$$
$$= \sum_j \beta_j^Z \text{bias}_j^Z \pi_j(1-\pi_j)$$
$$= \sum_j \beta_j^Z \pi_j(1-\pi_j)\Big(\beta_j^Z\big[2\theta_m^Z\theta_p^Z h_Z^2 + (\theta_m^Z\theta_p^Y + \theta_p^Z\theta_m^Y)\rho_{ZY}\big] + \beta_j^Z\beta_j^Y\big[(\theta_m^Y\theta_p^Z + \theta_p^Y\theta_m^Z)h_Z^2 + 2\theta_m^Y\theta_p^Y\rho_{ZY}\big]$$
$$+ \beta_j^Z\beta_j^{U_m}(\theta_p^Z h_Z^2 + \theta_p^Y\rho_{ZY}) + \beta_j^Z\beta_j^{U_p}(\theta_m^Z h_Z^2 + \theta_m^Y\rho_{ZY})\Big)$$
$$= h_Z^2\big[2\theta_m^Z\theta_p^Z h_Z^2 + (\theta_m^Z\theta_p^Y + \theta_p^Z\theta_m^Y)\rho_{ZY}\big] + \rho_{ZY}\big[(\theta_m^Y\theta_p^Z + \theta_p^Y\theta_m^Z)h_Z^2 + 2\theta_m^Y\theta_p^Y\rho_{ZY}\big]$$
$$= 2\big[\theta_m^Z\theta_p^Z h_Z^4 + \theta_m^Y\theta_p^Y\rho_{ZY}^2 + (\theta_m^Z\theta_p^Y + \theta_p^Z\theta_m^Y)h_Z^2\rho_{ZY}\big]. \tag{26}$$

By symmetry, the excess variance for the outcome trait is

$$\text{ExcessVariance}^Y = 2\big[\theta_m^Z\theta_p^Z\rho_{ZY}^2 + \theta_m^Y\theta_p^Y h_Y^4 + (\theta_m^Z\theta_p^Y + \theta_p^Z\theta_m^Y)h_Y^2\rho_{ZY}\big]. \tag{27}$$

The intuition for these expressions is simple. Genetic variance for $Z$ trait increases when parents sort on $Z$ (i.e., $\theta_m^Z \neq 0$ and $\theta_p^Z \neq 0$), when they sort on a genetically correlated trait (i.e., $\theta_m^Y \neq 0$, $\theta_p^Y \neq 0$, $\rho_{ZY}^2 \neq 0$)), and when there is cross trait sorting between a pair of genetically correlated traits. The parallel statements hold for trait $Y$.

Now considering the excess covariance,

$$\text{ExcessCovariance}^{ZY} = \text{Cov}\big(\textstyle\sum_j x_j\beta_j^Z, \sum_j x_j\beta_j^Y\big) - \sum_j \text{Cov}\big(x_j\beta_j^Z, x_j\beta_j^Y\big). \tag{28}$$

Following parallel steps to the calculation of the excess variance obtains

$$\text{ExcessCovariance}^{ZY} = 2\big(\theta_m^Z\theta_p^Z h_Z^2 + \theta_m^Y\theta_p^Y h_Y^2\big)\rho_{ZY} + \big(\theta_m^Z\theta_p^Y + \theta_p^Z\theta_m^Y\big)\big(h_Z^2 h_Y^2 + \rho_{ZY}^2\big) \tag{29}$$

As with the excess variance, the first term of this expression implies that excess covariance will increase if both parents sort on the treatment or outcome trait, and the second term implies that excess covariance increases if there is multi- or cross-trait sorting.

The formula for the excess genetic correlation does not have a clean closed form except in certain simple cases.

*AM-induced LD with arbitrary AM*

Above we have derived a simple expression of the effect of AM on AM-induced LD and GWAS bias after a single generation of sorting. We now consider the case where the parents potential come from a population that may have had non-random mating for several generations. We do not assume that the population is in equilibrium or that the parameters defining the sorting have been constant over time. However, by inductive reasoning, we will show that if, in the parents' generation, the covariance between a pair of SNPs is of the form

$$\mathrm{Cov}(x_j, x_k) = \left[C_{1,jk}\alpha_j^m\alpha_k^m + C_{2,jk}\alpha_j^m\alpha_k^p + C_{3,jk}\alpha_j^p\alpha_k^m + C_{4,jk}\alpha_j^p\alpha_k^p\right], \quad (30)$$

where $C_{1,jk}$, $C_{2,jk}$, $C_{3,jk}$, and $C_{4,jk}$ are some constants, then the covariance between those SNPs in the children's generation is also of that form (with potentially different constants). For example, in the case where the parents are born from a randomly mating population, $C_{1,jk} = C_{2,jk} = C_{3,jk} = C_{4,jk} = 0$ for the parents. In the children's generation, $C_{1,jk} = C_{4,jk} = 0$ and $C_{2,jk} = C_{3,jk} = \pi_j(1-\pi_j)\pi_k(1-\pi_k)$.

As before, we have

$$\begin{aligned}\mathrm{Cov}(x_j, x_k) &= \mathrm{Cov}(x_{m,j}^t + x_{p,j}^t, x_{m,k}^t + x_{p,k}^t) \\ &= \mathrm{Cov}(x_{m,j}^t, x_{m,k}^t) + \mathrm{Cov}(x_{m,j}^t, x_{p,k}^t) + \mathrm{Cov}(x_{p,j}^t, x_{m,k}^t) + \mathrm{Cov}(x_{p,j}^t, x_{p,k}^t)\end{aligned}$$

In this case where the parents come from a generation with AM, $\mathrm{Cov}(x_{m,j}^t, x_{m,k}^t)$ and $(x_{p,j}^t, x_{p,k}^t)$ are not equal to zero. Instead, assuming that recombination is equally likely between any pair of variants, we have

$$\mathrm{Cov}(x_{m,j}^t, x_{m,k}^t) = \mathrm{Cov}(x_{p,j}^t, x_{p,k}^t) = \frac{1}{4}\left[C_{1,jk}\alpha_j^m\alpha_k^m + C_{2,jk}\alpha_j^m\alpha_k^p + C_{3,jk}\alpha_j^p\alpha_k^m + C_{4,jk}\alpha_j^p\alpha_k^p\right]$$

The derivation of $\mathrm{Cov}(x_{m,j}^t, x_{p,k}^t)$ follows the same steps as above until we reach the step

$$\mathrm{Cov}(x_{m,j}^t, x_{p,k}^t) = \int \frac{f(S|x_{m,j}^t=1)f(S|x_{p,k}^t=1)}{f(S)}dS\,\pi_j\pi_k - \pi_j\pi_k. \quad (31)$$

In the AM case,

$$\begin{aligned}\mathrm{E}(S|x_{m,j}^t = 1) &= \mathrm{E}\left(\sum_\ell \alpha_\ell^m(x_{m,\ell}^t + x_{m,\ell}^u) + e_m|x_{m,j}^t = 1\right) \\ &= \alpha_j^m + \sum_{\ell\neq j}\alpha_\ell^m\mathrm{E}(x_{m,\ell}^t|x_{m,j}^t = 1) + \sum_\ell \alpha_\ell^m\mathrm{E}(x_{m,\ell}^u|x_{m,j}^t = 1) + \mathrm{E}(e_m|x_{m,j}^t = 1) \\ &= \alpha_j^m + \sum_{\ell\neq j}\alpha_\ell^m\mathrm{E}(x_{m,\ell}^u|x_{m,j}^t = 1) + \sum_\ell \alpha_\ell^m\mathrm{E}(x_{m,\ell}^u|x_{m,j}^t = 1) + \mathrm{E}(e_m|x_{m,j}^t = 1) \\ &= \alpha_j^m - \alpha_\ell^m\mathrm{E}(x_{m,j}^u|x_{m,j}^t = 1) + 2\textstyle\sum_\ell \alpha_\ell^m\mathrm{E}(x_{m,\ell}^u|x_{m,j}^t = 1) + \mathrm{E}(e_m|x_{m,j}^t = 1). \quad (32)\end{aligned}$$

Next, we note that by the properties of linear projections

$$\mathrm{E}(x_{m,k}^u|x_{m,j}^t = 1) = \left[\pi_k - \frac{\mathrm{Cov}(x_{m,j}^u, x_{m,j}^t)}{\mathrm{Cov}(x_{m,j}^t)}(1-\pi_j)\right]. \quad (33)$$

By substituting (33) into (32) and rearranging terms, we have

$$\mathrm{E}\left(S|x_{m,j}^{t}=1\right)=A_{m,j}\alpha_j^m\left(1-\pi_j\right)+A_{p,j}\alpha_j^p\left(1-\pi_j\right)+\mathrm{E}(S), \tag{34}$$

where

$$A_{m,j}=1+\frac{\mathrm{Cov}\left(x_{m,j}^{u},x_{m,j}^{t}\right)}{\mathrm{Cov}\left(x_{m,j}^{t}\right)}-\frac{1}{2}\sum_{\ell}\alpha_\ell^m\left[\frac{C_{1,j\ell}\alpha_\ell^m+C_{2,j\ell}\alpha_\ell^p}{\mathrm{Cov}\left(x_{m,j}^{t}\right)}\right] \tag{35}$$

$$A_{p,j}=-\frac{1}{2}\sum_{\ell}\alpha_\ell^m\left[\frac{C_{3,j\ell}\alpha_\ell^m+C_{4,j\ell}\alpha_\ell^p}{\mathrm{Cov}\left(x_{m,j}^{t}\right)}\right]. \tag{36}$$

Following a parallel derivation, we have

$$\mathrm{E}\left(S|x_{p,k}^{t}=1\right)=A_{p,k}\alpha_k^p(1-\pi_k)+A_{m,k}\alpha_k^m(1-\pi_k)+\mathrm{E}(S), \tag{37}$$

where

$$A_{p,k}=1+\frac{\mathrm{Cov}\left(x_{p,k}^{u},x_{p,k}^{t}\right)}{\mathrm{Cov}\left(x_{p,k}^{t}\right)}-\frac{1}{2}\sum_{\ell}\alpha_\ell^p\left[\frac{C_{1,k\ell}\alpha_\ell^p+C_{2,k\ell}\alpha_\ell^m}{\mathrm{Cov}\left(x_{p,k}^{t}\right)}\right] \tag{38}$$

$$A_{m,k}=-\frac{1}{2}\sum_{\ell}\alpha_\ell^p\left[\frac{C_{3,k\ell}\alpha_\ell^p+C_{4,k\ell}\alpha_\ell^m}{\mathrm{Cov}\left(x_{m,j}^{t}\right)}\right]. \tag{39}$$

As before, substituting (34) and (37) into (31), completing the square, and rearranging terms gives us

$$\begin{aligned}\mathrm{Cov}\left(x_{m,j}^{t},x_{p,k}^{t}\right)&\approx\int\sqrt{2\pi}e^{-\frac{1}{2}\left[\left(S-E\left(S|x_{m,j}^{t}=1\right)\right)^2+\left(S-E\left(S|x_{p,k}^{t}=1\right)\right)^2-\left(S-E(S)\right)^2\right]}dS\,\pi_j\pi_k-\pi_j\pi_k\\&=e^{\left(\alpha_j^mA_{m,j}+\alpha_j^pA_{p,j}\right)\left(\alpha_k^mA_{m,k}+\alpha_k^pA_{p,k}\right)(1-\pi_j)(1-\pi_k)}\,\pi_j\pi_k-\pi_j\pi_k\\&\approx\left(\alpha_j^mA_{m,j}+\alpha_j^pA_{p,j}\right)\left(\alpha_k^mA_{m,k}+\alpha_k^pA_{p,k}\right)\pi_j\left(1-\pi_j\right)\pi_k(1-\pi_k)\end{aligned} \tag{40}$$

And by symmetry, we have

$$\mathrm{Cov}\left(x_{m,k}^{t},x_{p,j}^{t}\right)\approx\left(\alpha_k^mB_{m,k}+\alpha_k^pB_{p,k}\right)\left(\alpha_j^mB_{m,j}+\alpha_j^pB_{p,j}\right)\pi_j\left(1-\pi_j\right)\pi_k(1-\pi_k) \tag{41}$$

with

$$B_{m,k}=1+\frac{\mathrm{Cov}(x_{m,k}^{u},x_{m,k}^{t})}{\mathrm{Cov}(x_{m,k}^{t})}-\frac{1}{2}\sum_{\ell}\alpha_\ell^m\left[\frac{C_{1,k\ell}\alpha_\ell^m+C_{2,k\ell}\alpha_\ell^p}{\mathrm{Cov}(x_{m,k}^{t})}\right] \tag{42}$$

$$B_{p,k}=-\frac{1}{2}\sum_{\ell}\alpha_\ell^m\left[\frac{C_{3,k\ell}\alpha_\ell^m+C_{4,k\ell}\alpha_\ell^p}{\mathrm{Cov}(x_{m,k}^{t})}\right]. \tag{43}$$

$$B_{p,j} = 1 + \frac{\mathrm{Cov}\left(x^u_{p,j}, x^t_{p,j}\right)}{\mathrm{Cov}\left(x^t_{p,j}\right)} - \frac{1}{2}\sum_\ell \alpha^p_\ell \left[\frac{C_{1,j\ell}\alpha^p_\ell + C_{2,j\ell}\alpha^m_\ell}{\mathrm{Cov}\left(x^t_{p,j}\right)}\right] \tag{44}$$

$$B_{m,j} = -\frac{1}{2}\sum_\ell \alpha^p_\ell \left[\frac{C_{3,j\ell}\alpha^p_\ell + C_{4,j\ell}\alpha^m_\ell}{\mathrm{Cov}\left(x^t_{m,j}\right)}\right]. \tag{45}$$

Putting all this together gives us

$$\begin{aligned}
\mathrm{Cov}(x_j, x_k) &= \left(\alpha^m_j A_{m,j} + \alpha^p_j A_{p,j}\right)\left(\alpha^m_k A_{m,k} + \alpha^p_k A_{p,k}\right)\pi_j(1-\pi_j)\pi_k(1-\pi_k) \\
&\quad + \left(\alpha^m_k B_{m,k} + \alpha^p_k B_{p,k}\right)\left(\alpha^m_j B_{m,j} + \alpha^p_j B_{p,j}\right)\pi_j(1-\pi_j)\pi_k(1-\pi_k) \\
&\quad + \frac{1}{2}\left[C_{1,jk}\alpha^m_j\alpha^m_k + C_{2,jk}\alpha^m_j\alpha^p_k + C_{3,jk}\alpha^p_j\alpha^m_k + C_{4,jk}\alpha^p_j\alpha^p_k\right] \\
&= \alpha^m_j\alpha^m_k\left(\left[A_{m,j}A_{m,k} + B_{m,j}B_{m,k}\right]\pi_j(1-\pi_j)\pi_k(1-\pi_k) + \frac{1}{2}C_{1,jk}\right) \\
&\quad + \alpha^m_j\alpha^p_k\left(\left[A_{m,j}A_{p,k} + B_{m,j}B_{p,k}\right]\pi_j(1-\pi_j)\pi_k(1-\pi_k) + \frac{1}{2}C_{2,jk}\right) \\
&\quad + \alpha^p_j\alpha^m_k\left(\left[A_{p,j}A_{m,k} + B_{p,j}B_{m,k}\right]\pi_j(1-\pi_j)\pi_k(1-\pi_k) + \frac{1}{2}C_{3,jk}\right) \\
&\quad + \alpha^p_j\alpha^p_k\left(\left[A_{p,j}A_{p,k} + B_{p,j}B_{p,k}\right]\pi_j(1-\pi_j)\pi_k(1-\pi_k) + \frac{1}{2}C_{4,jk}\right).
\end{aligned}$$

This expression is of the form

$$\left[C_{1,jk}\alpha^m_j\alpha^m_k + C_{2,jk}\alpha^m_j\alpha^p_k + C_{3,jk}\alpha^p_j\alpha^m_k + C_{4,jk}\alpha^p_j\alpha^p_k\right].$$

So by induction, if the covariance between pairs of SNPs is of this form in one generation (as it is in a randomly mating population), it must be of the same form in any subsequent generation under our model of sorting.

*GWAS Bias with arbitrary AM*

As before, we would like to evaluate the bias of GWAS estimates for the treatment trait under a general model of sorting. The initial steps in this case are identical to the first-generation of AM derivation such that

$$\mathrm{Bias}^Z_j = \frac{\sum_{k\neq j}\beta^Z_k \mathrm{Cov}(x_j, x_k)}{\mathrm{Var}(x_j)}. \tag{46}$$

Here, we substitute in the general form of AM-induced LD, eq. (30), into (46), giving us

$$\begin{aligned}
\mathrm{Bias}^Z_j &= \frac{\sum_{k\neq j}\beta^Z_k\left[C_{1,jk}\alpha^m_j\alpha^m_k + C_{2,jk}\alpha^m_j\alpha^p_k + C_{3,jk}\alpha^p_j\alpha^m_k + C_{4,jk}\alpha^p_j\alpha^p_k\right]}{\mathrm{Var}(x_j)}. \\
&= \alpha^m_j\left(\frac{\sum_{k\neq j}\beta^Z_k C_{1,jk}\alpha^m_k}{\mathrm{Var}(x_j)} + \frac{\sum_{k\neq j}\beta^Z_k C_{2,jk}\alpha^p_k}{\mathrm{Var}(x_j)}\right) + \alpha^p_j\left(\frac{\sum_{k\neq j}\beta^Z_k C_{3,jk}\alpha^m_k}{\mathrm{Var}(x_j)} + \frac{\sum_{k\neq j}\beta^Z_k C_{4,jk}\alpha^p_k}{\mathrm{Var}(x_j)}\right)
\end{aligned} \tag{47}$$

Next, substituting in (14-15), we obtain

$$\begin{aligned}\text{Bias}_j^Z &= \left(\theta_m^Z\beta_j^Z + \theta_m^Y\beta_j^Y + \beta_j^{U_m}\right)\left(\frac{\sum_{k\neq j}\beta_k^Z C_{1,jk}\alpha_k^m}{\text{Var}(x_j)} + \frac{\sum_{k\neq j}\beta_k^Z C_{2,jk}\alpha_k^p}{\text{Var}(x_j)}\right)\\ &\quad + \left(\theta_p^Z\beta_j^Z + \theta_p^Y\beta_j^Y + \beta_j^{U_p}\right)\left(\frac{\sum_{k\neq j}\beta_k^Z C_{3,jk}\alpha_k^m}{\text{Var}(x_j)} + \frac{\sum_{k\neq j}\beta_k^Z C_{4,jk}\alpha_k^p}{\text{Var}(x_j)}\right)\\ &= \beta_j^Z\left[\theta_m^Z\left(\frac{\sum_{k\neq j}\beta_k^Z C_{1,jk}\alpha_k^m}{\text{Var}(x_j)} + \frac{\sum_{k\neq j}\beta_k^Z C_{2,jk}\alpha_k^p}{\text{Var}(x_j)}\right) + \theta_p^Z\left(\frac{\sum_{k\neq j}\beta_k^Z C_{3,jk}\alpha_k^m}{\text{Var}(x_j)} + \frac{\sum_{k\neq j}\beta_k^Z C_{4,jk}\alpha_k^p}{\text{Var}(x_j)}\right)\right]\\ &\quad +\beta_j^Y\left[\theta_m^Y\left(\frac{\sum_{k\neq j}\beta_k^Z C_{1,jk}\alpha_k^m}{\text{Var}(x_j)} + \frac{\sum_{k\neq j}\beta_k^Z C_{2,jk}\alpha_k^p}{\text{Var}(x_j)}\right) + \theta_p^Y\left(\frac{\sum_{k\neq j}\beta_k^Z C_{3,jk}\alpha_k^m}{\text{Var}(x_j)} + \frac{\sum_{k\neq j}\beta_k^Z C_{4,jk}\alpha_k^p}{\text{Var}(x_j)}\right)\right]\\ &\quad + \beta_j^{U_m}\left(\frac{\sum_{k\neq j}\beta_k^Z C_{1,jk}\alpha_k^m}{\text{Var}(x_j)} + \frac{\sum_{k\neq j}\beta_k^Z C_{2,jk}\alpha_k^p}{\text{Var}(x_j)}\right)\\ &\quad + \beta_j^{U_p}\left(\frac{\sum_{k\neq j}\beta_k^Z C_{3,jk}\alpha_k^m}{\text{Var}(x_j)} + \frac{\alpha_j^Z\sum_{k\neq j}\beta_k^Z C_{4,jk}\alpha_k^p}{\text{Var}(x_j)}\right).\end{aligned}$$

Notice that this is linear in $\beta_j^Z$, $\beta_j^Y$, $\beta_j^{U_m}$, and $\beta_j^{U_p}$. Therefore, we see that in any generation of arbitrary sorting, the bias in GWAS estimates for the treatment trait will always be linear in the true conditional associations of SNPs on the treatment trait.

This property of the bias is useful because there is risk of SNP-selection bias if $\partial Bias_j^Z/\partial\beta_j^Y = 0$. By the linearity above, this derivative is a constant, making it easier to determine whether the condition holds.

### II. Simulations

*Simulation design.*

Studies already exist with large-scale simulations evaluating the effect of AM on genetic correlation (Border et al., 2022) and MR analyses (Hartwig et al., 2018). Here, we consider a series of simplified simulations that confirm some of their points, fill in the gaps that were missed in their analyses, and provide counter examples to some of their broader claims. We also evaluate the performance of our adjustments to genetic correlation and MR estimates under various conditions.

In these simulations, we consider four genetic architectures: (1) no correlation/perfect pleiotropy, (2) partial correlation/perfect pleiotropy (3) no pleiotropy, (4) partial pleiotropy. Under (1), the simulated effect of each SNP on each trait is drawn from a bivariate standard normal distribution with no correlation. Under (2), simulated effect of each SNP is drawn from a bivariate normal with mean zero, variance one, and a correlation of 0.5. Under (3), half of SNPs have simulated effects drawn from standard normal for trait 1 and have an effect of zero on trait 2, and for the residual half, the SNPs have a simulated effect of zero on trait 1 and an effect drawn from a standard normal distribution for trait 2. Under (4), 25% of SNPs affect trait 1 but not trait 2, 25% affect trait 2 but not trait 1, and 50% affect both traits. In this partial pleiotropy case, effect sizes are drawn from a standard normal distribution, with equal effects on trait 1 and 2 for pleiotropic SNPs. See Figure S1 for illustrations of these genetic architectures.

In the MR analyses, each of these approaches can be mapped to different ways that the exclusion restriction may be met or violated. For example, cases (1)-(3) correspond to scenarios where no SNP satisfies the exclusion restriction. Case (4) corresponds to a case where half of SNPs do. Notably, the 25% of SNPs that have no effect on the treatment trait will generally be excluded from MR since they would not meet the statistical significance criterion for inclusion. As a result, more than half of included SNPs would satisfy exclusion and therefore (in the randomly mating population) the assumptions of median-based MR (and Mode-based MR) should be met. Despite this, if we assume that the causal effect of the treatment on the outcome is 0 and 0.5 for cases (1) and (2), respectively, both median-based and mode-based MR will still be consistent in a randomly mating population since the median and modal SNP satisfy exclusion.

We simulate 4 sorting mechanisms: (1) sorting on just the treatment phenotype, (2) sorting on just the outcome phenotype, (3) cross-trait sorting, and (4) multi-trait sorting. Cross-trait sorting is when males with a high value for trait 1 are more likely to mate with females with a high value of trait 2 (independent of their value for trait 1). When considering the effect of AM on genetic variance or genetic correlation, due to their symmetry, we collapse mechanisms (1) and (2) into a single mechanism, which we call single-trait sorting. Multi-trait sorting is when people sort based on linear combination of both trait 1 and trait 2. In our multi-trait sorting simulation, individuals sort on the linear combination of both traits with a weight of 0.8 on trait 1 and 0.2 on trait 2. In each simulation, in the first generation of 10,000 people, we simulate a population that is assumed to be randomly mating. (That is, each SNP is uncorrelated with each other SNP in the genome.) To maximize statistical power and to increase the effect of AM on AM-induced LD in subsequent generations, we simulate both the treatment and outcome traits with a heritability of one. To produce the next generation, we randomly assign half of the population as male and the other half as female. We then rank-match one person from each group to the corresponding person from the other group. (E.g., in the single-trait sorting cases, the male with the highest trait value would be matched with the female with the highest trait value.) Because our simulated sample is so large, this results in a near-perfect correlation of the traits on which mates are sorting. We model near-perfect sorting for computational reasons since stronger sorting will give rise to large amounts of AM-induced LD in few generations.

Each pair is simulated to have two children (which maintains the population size). For simplicity, we assume perfect recombination between each pair of SNPs. This matching and mating procedure is repeated until we have four generations of data, with the earliest generation of data corresponding to random mating and the fourth generation having the largest impacts of AM. We then calculate the genetic correlation of the two traits for each generation and the MR estimate of the effect of the treatment on the outcome, each with and without the proposed AM adjustment. We also calculate the increase in the genetic variance of each trait relative to the genetic variance in the first (randomly mating) generation.

The PGIs used for the AM adjustments are simulated in the following way. To account for the potential of causal SNPs that are not included in the PGI, we simulate cases where all causal SNPs are included in the PGI and when only every other SNP is included. To account for estimation error in the PGI weights, the weight for SNP $j$ and trait $Z$ was set as

$$w_j = \phi E_j^Z + (1 - \phi)\beta_j^Z \tag{48}$$

where $\phi \in \{0.0, 0.2, 0.4, 0.6, 0.8, 1.0\}$ is the parameter that sets the fraction of the PGI that is due to error, $E_j^Z \sim N(0,1)$ is an error term, and $\beta_j^Z$ is the simulated effect size (also drawn from a standard normal distribution). Combing both sources of error, this means we produce 12 PGIs for each replication of the simulation.

The parameter $\phi$ in this paper and the parameter $\rho$ defined in Becker et al. (2021) are related. Becker et al. defines $\rho$ as the amount of shrinkage observed in a regression of some trait on the PGI for that trait. Under our model

$$\phi = 1 - \frac{1}{\rho^2}.$$

Becker et al. estimate $\rho = 1.3$ for educational attainment, which implies that $\phi = .4$ in that case.

*Simulation study results*

Supplementary Figure S4 reports our simulation results related to how AM inflates genetic variance and Supplementary Figure S5 reports the results for genetic correlation using a PGI with no error as a best-case scenario (see Supplementary Tables S1 and S3 for complete results). The simulations have two main implications for the effect on AM on genetic variance. First, single-trait AM on trait 1 has no effect on the genetic variance of trait 2 if the traits are uncorrelated in the randomly mating population (i.e., the $r_g$ = 0 and No Pleiotropy cases). Second, we see inflation of the genetic variance for both traits in all other settings, though this inflation is very small in the multi-trait AM cases for the uncorrelated genetic architectures. Both of these observations are consistent with the theory summarized in Main Text Table 1.

The simulations also have two main implications on the effect of AM on genetic correlation. First, single-trait AM inflates the genetic correlation between pairs of traits when there is a positive genetic correlation in the randomly mating population, but it does not affect the genetic correlation when the traits are not initially genetically correlated. This is consistent with the theoretical results of Gianola (1982) based on a bivariate normal model. Our simulations show that his conclusions hold also under our more complicated partial pleiotropy architecture. Second, cross-trait and multi-trait AM always inflate genetic correlation, a replication of the results reported in Border et al. (2022). When we apply our AM adjustment using a PGI without error to our simulated data, we see that we mostly recover the random mating estimate of genetic correlation, as expected based on our theory. The partial recovery is due to the fact our adjustment does not correct for any within-chromosome effects on LD. As expected, however, the residual inflation after the adjustment is generally small. When we consider calculate the adjusted genetic correlation in a randomly mating population, the adjustment is not biased on average, though the adjustment does become unstable for large values of $\phi$ (i.e., very noisy PGIs). In AM populations, however, some downward bias is introduced, implying that our approach over-adjusts for AM. The implication of this over adjustment is that, our method will not identify a role of AM in genetic correlation estimates when there is none; however, if our method estimates a non-zero effect of AM, that estimate will be an upper bound on the true effect of AM on the genetic correlation.

The MR simulation results presented in Supplementary Figure S7 and Supplementary Tables S2 and S4 produce several important conclusions. First, for the architectures considered, single-trait AM on the treatment trait generally does not influence MR estimates except in the partial pleiotropy case, where it produces a bias toward the null. The reason for this bias in the partial pleiotropy case is that AM produces LD between the SNPs that are causal for both traits and the SNPs that are casual for only the "treatment" trait. Because MR analyses rely on pre-selection of instrument based on GWAS summary statistics, this AM-induced LD will result in preferential selection of SNP-treatment associations with inflated coefficients (is this a type of winner's curse? If so, I think we should use that phrase). This AM-induced LD will not inflate the outcome GWAS coefficients because those SNPs do not affect the outcome trait. Because the treatment GWAS coefficient is in the denominator of the MR ratios, this will result in bias towards the null. This attenuation does not affect the $r_g$=.5 case because there are no SNPs that affect the treatment trait and not the outcome trait.

Second, MR estimates are inflated away from zero when there is sorting on the outcome trait. The mechanism by which this occurs is analogous to the reason that estimates are biased toward the null when there is sorting on the treatment.

Third, cross-trait and multi-trait AM inflates MR estimates in all cases except the partial pleiotropy case. This is because cross- and multi-trait AM generally increase LD between SNPs if one increases the outcome trait and the other increases the treatment trait. This will push the GWAS effects (which will capture the effects of both SNPs in the sorting population) to have concordant associations. In the case of the partial pleiotropy genetic architecture, cross-trait assortative mating inflates both the treatment and outcome GWAS associations equally since the fraction of SNPs that affect one trait and not the other are equally sized. Therefore, the inflation of the associations cancels out in the MR ratios. In the multi-trait sorting case, because there is greater weight on the treatment trait, the inflation of the GWAS associations is larger, which actually leads to a downward bias of the MR associations.

Fourth, in all cases, if a PGI without error is used, we see that our adjusted MR estimates is closer to the MR estimate that would be obtained under random mating than the unadjusted estimates. A small amount of the residual bias is due to within-chromosome LD, though the remaining bias is due to other issues discussed in the next paragraph.

Fifth, while the AM-adjusted MR estimates mostly obtain the random-mating MR estimate, large gaps remain in the Outcome AM case when $r_g$=.5 and in the Cross- and Multi-trait AM cases for both perfect-pleiotropy architectures. This is due to SNP-selection bias. In these cases, the GWAS associations are inflated more by AM for SNPs that have large, sign-concordant effects on both traits. Since such SNPs have especially inflated treatment associations, these SNPs are more likely to satisfy the significance criterion for inclusion in the MR study. For example, in the cross-trait AM/$r_g$=0 simulation, SNPs with concordant-signed effects and discordant-signed effects are equally likely to be included in the MR analysis in a randomly mating population (concordant: 8.7%, SE = 0.2%; discordant: 8.4%, SE = 0.2%), but in the fourth generation, concordant-signed SNPs have a 23.5% chance (SE = 0.2%) of being included compared to discordant-signed SNPs, which have only a 7.8% chance (SE = 0.2%). This bias is in addition to the bias from AM-induced LD and therefore is not accounted for by MR-adj. Because of this, we see that in many cases, over half of the bias in our MR study simulations appears to be due to selection of instruments.

Sixth, considering the results in Supplementary Table S4, error in the PGI can generate adjusted estimates that are downward biased, even in a randomly mating population. This is because the adjustment factor is the ratio of two very small numbers, so even a small amount of error can make the adjustment factor approximate a Cauchy distribution as the ratio of two mean-zero normal distributions. This implies that, unless very large samples are available to produce high powered PGIs, this approach may be impractical to adjust MR estimates.

Overall, our simulation results highlight that both single-trait, cross-trait, and multi-trait AM can have dramatic effects on the interpretation of genetic correlation and MR studies. Our adjustment can be a useful tool for understanding the role of AM on genetic correlation estimates, but it is less useful for MR applications due to SNP-selection bias and bias due to noisy in the PGI.

**III. Mendelian randomization Estimators**

In this paper, we consider three different Mendelian randomization estimators: MR-Egger (Bowden et al. 2015), MR-median (Bowden et al. 2016), and MR-Mode (Guo et al., 2018). Each of these approaches rely on different assumptions about the SNPs used in the analysis. A more detailed summary of each of them has been described elsewhere (Sanderson et al., 2022), but we here we provide a simple, high-level description of each of them below. We also provide more detailed information about how they were implemented in the analyses contained in this paper.

*Inverse-Variance Weighted MR (MR-IVW)*

Before describe the three methods used in this paper, we describe what might be considered the classic estimator: inverse-variance weighted (IVW) MR. We do not use this estimator in our analyses since it relies on strictly stronger assumptions that the other three estimators considered. For this estimator, the MR ratios ($\beta_j^Y/\beta_j^Z$) are first calculated for each SNP. Then, a weighted average is taken across all the ratios, where the weight for SNP $j$, $w_j$, is the inverse variance of the treatment GWAS association, $w_j = 1/SE_j^2$.

The usual assumption for this estimator is that all included SNPs satisfy the exclusion restriction. Under random mating this condition is generally met by the “no pleiotropy” architecture of our simulations since the SNPs that violate exclusion (i.e., those which do not affect the treatment but do affect the outcome) are likely to be omitted during the selection step. The estimator will also be consistent under the knife-edge case that all violations average out to be zero. Under random mating, this condition holds under the simulated full-pleiotropy architectures if we assume that the true causal effect is equal to $r_g$, and that deviations in the

MR ratio are due to exclusion violations of the exclusion restriction (which cancel out). However, in both cases, AM leads to violations of exclusion even in these special cases.

*MR-Egger*
In MR-Egger, the outcome GWAS associations are regressed onto the treatment GWAS associations with an intercept. The intercept in this model allows for biases in the MR ratios due to violations of the exclusion restriction that are not mean zero, as long as the bias term is uncorrelated with the treatment trait GWAS association. This assumption is called the Instrument Strength Independent of Direct Effect (InSIDE) assumption.

Because the INSIDE assumption is weaker than the assumption of MR-IVW, it is a consistent estimator in all cases where MR-IVW is consistent, including the no-pleiotropy and full-pleiotropy cases considered in our simulations. However, because the bias from AM in the treatment and outcome GWAS associations can be a function of the panmictic treatment trait GWAS associations, this means that AM will often cause the INSIDE assumption to fail under AM, creating bias in MR-Egger estimates.

*MR-Median*
For the MR-Median estimator, the MR ratios are calculated and the median of these ratios is used the estimated effect of the treatment on the outcome. Under random mating, the MR-Median estimator will be a consistent estimator of the causal effect if at least half of the included SNPs satisfy exclusion. This will hold in both the simulated no-pleiotropy and partial-pleiotropy cases. MR-Median can also be consistent when more than half of the included SNPs violate exclusion as long as less than half of the SNPs have positive bias due to exclusion violations and less than half of SNPs have negative bias. This holds in the full pleiotropy cases. Therefore, we expect MR-Median to be consistent for all simulated genetic architectures under random mating. However, this will hold for none of the genetic architectures under AM except in knife-edge cases where the bias affects the MR ratios in a balanced way.

*MR-Mode*
For the MR-Mode estimator, the MR ratios are calculated, and a distribution is fit to the ratios using kernel regression, a nonparametric distribution approach. We use a rectangular kernel with fixed bandwidth for computational simplicity. The mode of the estimated distribution is used as the MR estimate.

The intuitive explanation of the key assumption of MR-Mode is that a “plurality” of SNPs must satisfy exclusion; that is, more SNPs should satisfy exclusion, and therefore have a correct MR ratio, than the set of SNPs that have any other “incorrect” MR ratio. If this is the case, the mode will be a consistent estimator of the true causal effect. Under random mating, all of the simulated genetic architectures satisfy this criterion. However, under AM, this assumption will fail except in certain knife-edge scenarios.

While the MR-Mode estimator relies on the weakest assumptions of all the methods considered, it is also the least powerful. As a result, the simulation and empirical results presented in the paper, we focus on the MR-Median results, which are sufficiently robust under random mating for all scenarios and which are greater powered.

### IV. Empirical results for the effect of education on health

Supplementary Figure S9 displays the unadjusted and adjusted median-based MR estimates for the effect of education on health. The unadjusted MR-Egger and mode-based MR estimates follow a similar qualitative pattern to the unadjusted median-based estimates, though the mode-based estimates are estimated with much less precession (see Supplementary Table S7). These results highlight several main points. First, prior to the AM adjustment, the median MR estimates are statistically significant for six of nine considered traits (BMI, ever smoke, depressive symptoms, height, neuroticism, and self-rated health). Second, for each of these health outcomes, the AM adjustment reduces the magnitude of the estimates by an average of 30%. Third, each of these adjusted MR estimates remains statistically significant after the adjustment.

However, these adjusted results are potentially unreliable for two reasons. First, our adjustment only adjusts for AM-induced LD but not for SNP-selection bias, which can be as large as the bias from AM-induced LD. Second, as shown in our simulations, missing SNPs from the PGI and estimation error in the SNP effect-size estimates can lead to large, unpredictable biases in adjusted MR estimates. These issues can be seen most clearly in the example of the estimated “effect” of education on adult height. Our adjustment only reduces the estimated effect by 41%, though we would anticipate that the actual causal effect is likely negligible in a modern UK population like that of the UK Biobank (Minică et al., 2020). The residual 41% could be due to SNP-selection bias or error in the PGIs.
