## Supplementary Figures for "A General Approach to Adjusting Genetic Studies for Assortative Mating"

**Figure S1: Joint genetic architectures of simulated traits**

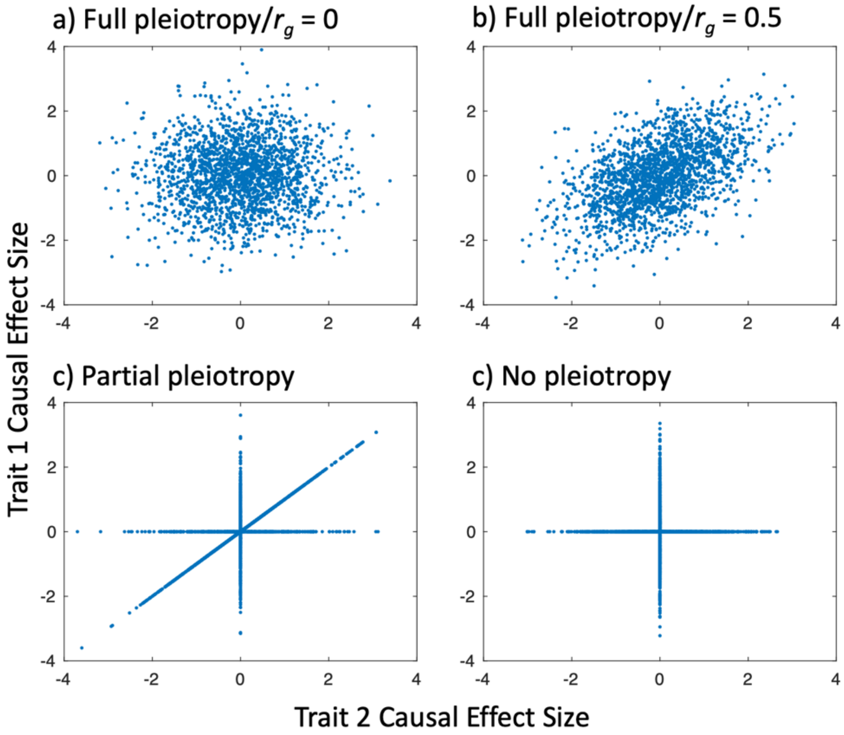


*Note*: Illustrations of the four genetic architectures modeled in our simulations. In **panel (a)**, effect sizes are drawn from a bivariate normal distribution with a mean of zero, a variance of one, and a correlation of zero. In **panel (b)**, effect sizes are drawn from a bivariate normal distribution with a mean of zero, a variance of one, and a correlation of 0.5. In **panel (c)**, for 50% of SNPs, effect sizes identical for both traits and drawn from a standard normal distribution. For 25% of SNPs, effects sizes are zero for trait 1 and drawn from a standard normal for trait 2. For the remain 25% of SNPs, effect sizes are drawn from a standard normal for trait 1 and are zero for trait 2. In **panel (d)**, for 50% of SNPs, effects sizes are zero for trait 1 and drawn from a standard normal for trait 2; and for the remain 50%, effect sizes are drawn from a standard normal for trait 1 and are zero for trait 2.

**Figure S2: Long-range LD Bias of GWAS Associations, $r_g$ = 0.0**

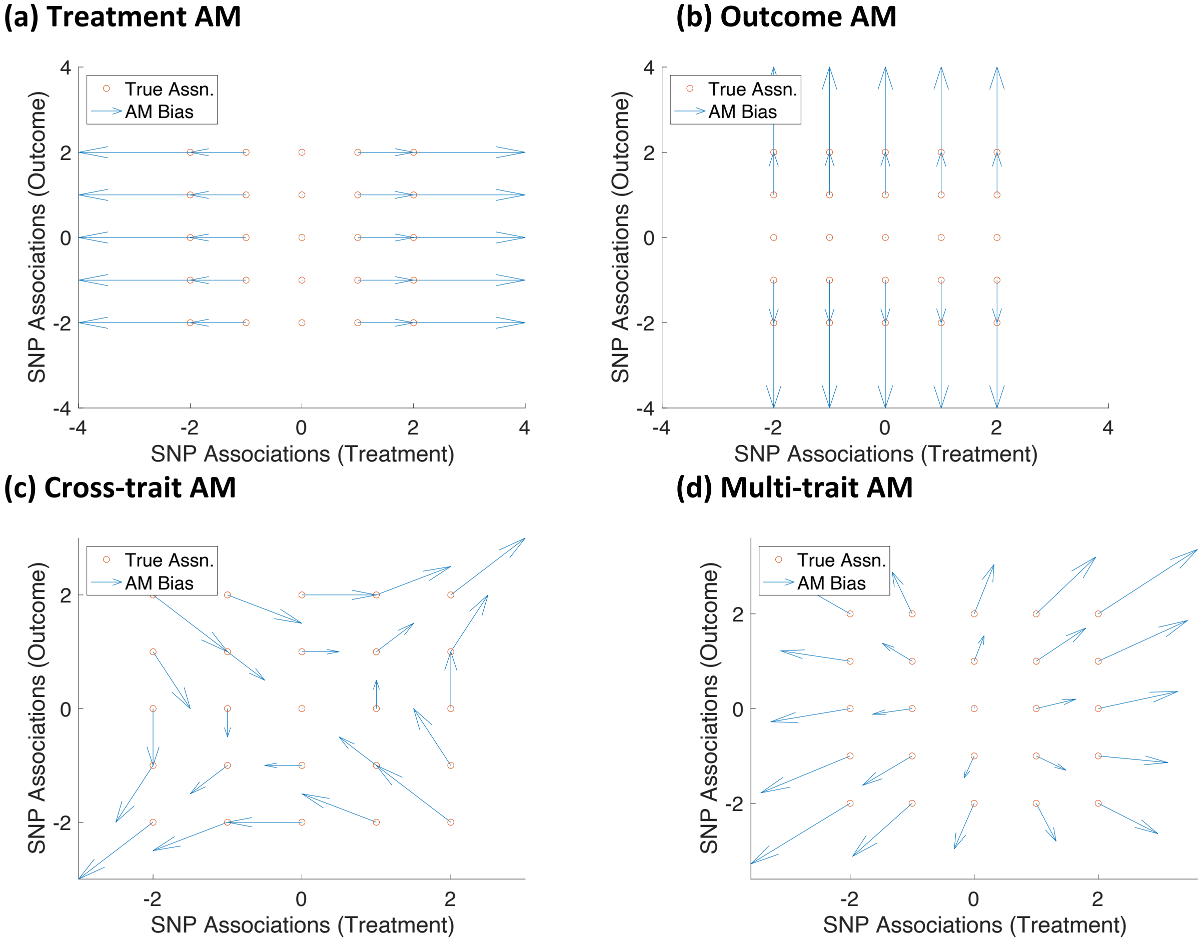


*Note*: This figure illustrates how different forms of AM bias GWAS associations for a pair of perfectly heritable traits when the genetic correlation between them is 0.0. In the multi-trait AM case, people sort on a linear combination of both traits with a weight of 0.8 on trait 1 and 0.2 on trait 2. The horizontal and vertical position of the red circles represent the true (i.e., panmictic) associations of a selected set of SNPs on the treatment and outcome traits, respectively. The blue arrows represent the how each of the SNP associations are affected due to direct sorting on the outcome based on the theory derived in the Supplementary Note.

**Figure S3: Long-range LD Bias of GWAS Associations, $r_g$ = 0.5**

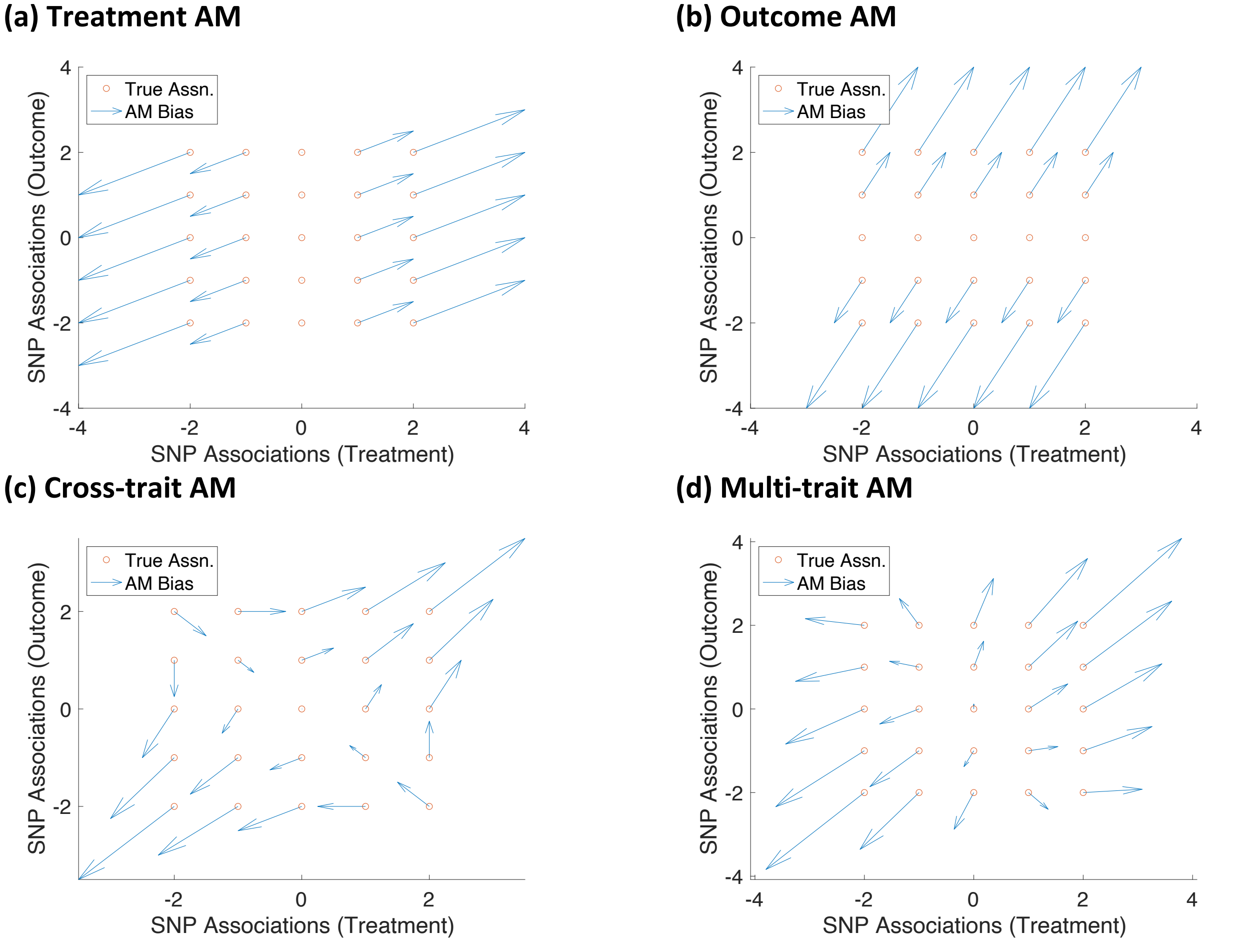


*Note*: This figure illustrates how different forms of AM bias GWAS associations for a pair of perfectly heritable traits when the genetic correlation between them is 0.5. In the multi-trait AM case, people sort on a linear combination of both traits with a weight of 0.8 on trait 1 and 0.2 on trait 2. The horizontal and vertical position of the red circles represent the true (i.e., panmictic) associations of a selected set of SNPs on the treatment and outcome traits, respectively. The blue arrows represent the how each of the SNP associations are affected due to direct sorting on the outcome based on the theory derived in the Supplementary Note.

**Figure S4: Simulated Effect of AM on Genetic Variance in Fourth Generation**

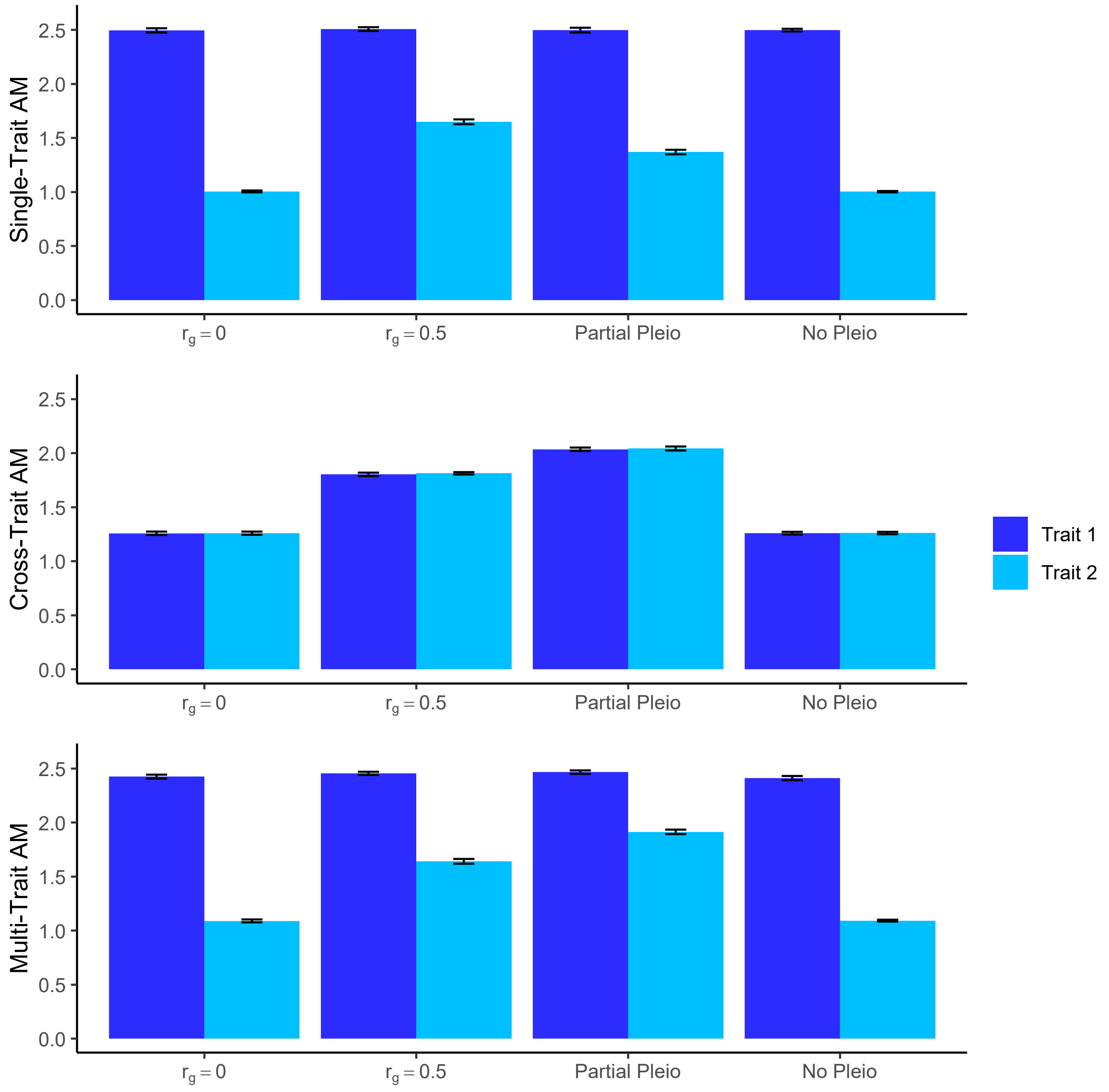


*Note*: This figure shows the simulations results for the effect of single-trait AM on trait 1, cross-trait AM, and multi-trait AM on the inflation of the genetic variance of each trait. In the multi-trait AM case, people sort on a linear combination of both traits with a weight of 0.8 on trait 1 and 0.2 on trait 2. We calculate the inflation of genetic variance in each replication as the ratio of the genetic variance in generation 4 over the genetic variance in generation 1, such that a value of one implies no inflation. We consider four different genetic architectures: total pleiotropy/zero correlation (**$r_g$=0**), total pleiotropy/positive correlation (**$r_g$=.5**), partial pleiotropy (**Part Pleio**), and no pleiotropy (**No Pleio**). In each replication of the simulation, we simulate a genome with 2000 SNPs even distributed across 20 chromosomes for 10,000 people. We perform 20 replications. The dark blue bars show the mean inflation for trait 1 and the light blue bars show the mean inflation for trait 2 over the 20 replications. The error bars correspond to the 95% confidence interval for the mean genetic correlation inflation.

**Figure S5: Simulated Effect of AM on Genetic Correlations**

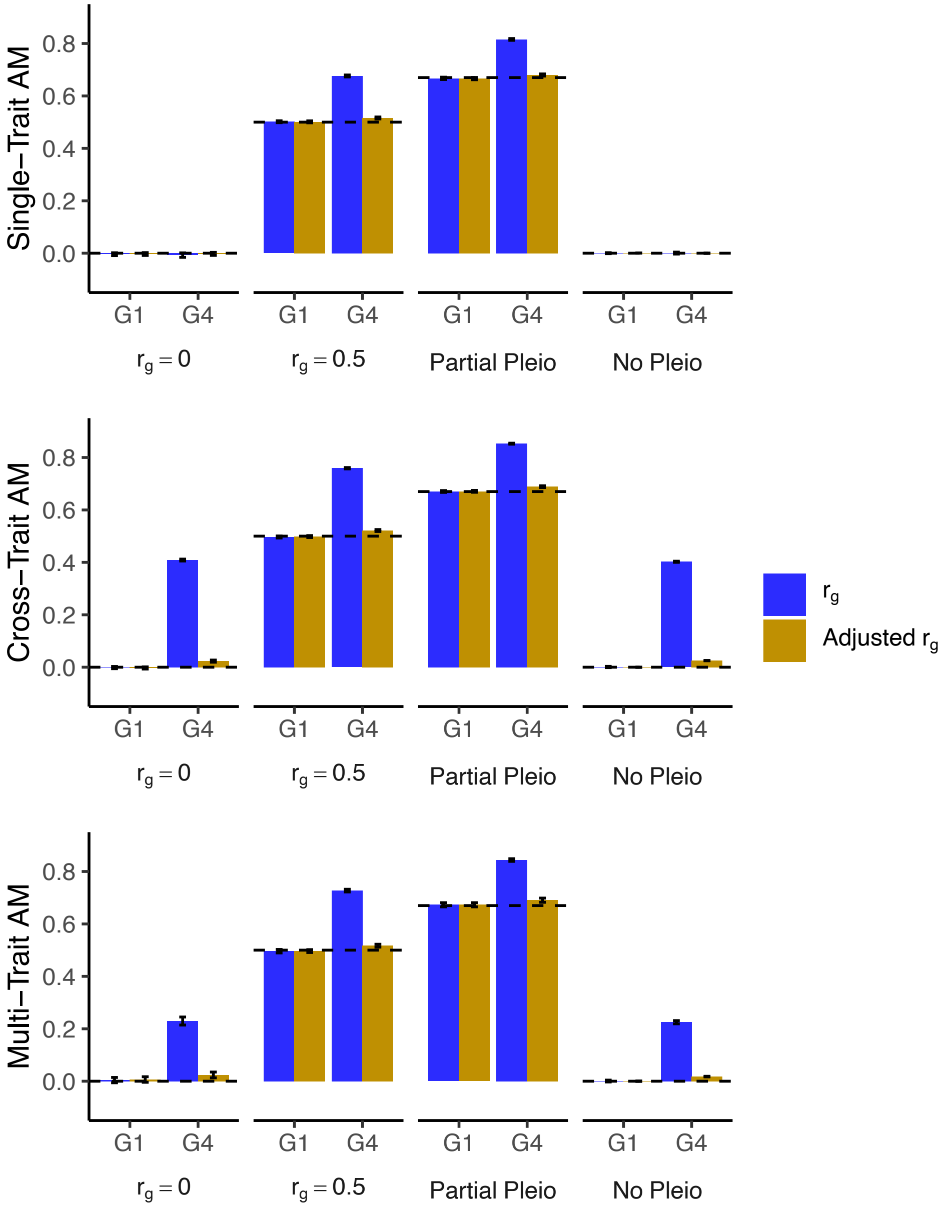


*Note*: This figure shows the simulations results for the effect of single-trait AM on trait 1, cross-trait AM, and multi-trait AM on genetic correlation (***r**<sub>g</sub>*). In the multi-trait AM case, people sort on a linear combination of both traits with a weight of 0.8 on trait 1 and 0.2 on trait 2. We consider four different genetic architectures: total pleiotropy/zero correlation (**$r_g$=0**), total pleiotropy/positive correlation (**$r_g$=.5**), partial pleiotropy (**Part Pleio**), and no pleiotropy (**No Pleio**). We evaluate the genetic correlation in the first generation (**G1**), which is modeled under a random mating assumption, and in the fourth generation (**G4**), which corresponds to a population undergoing three generations of AM. We model both single-trait and cross-trait AM. In each replication of the simulation, we simulate a genome with 2000 SNPs even distributed across 20 chromosomes for 10,000 people. We perform 20 replications. The blue bars show the mean unadjusted genetic correlation and the gold bars show the mean AM-adjusted genetic correlation over the 20 replications. The PGIs used in the adjustment are simulated with no error. The dotted line above each set of bars corresponds to the theoretical genetic correlation in a randomly mating population. The error bars correspond to the 95% confidence interval for the mean genetic correlation.

**Supplementary Figure S6: Summary of Error in AM Adjusted Genetic Correlation**

**(a)** Generation 2 **(b)** Generation 3

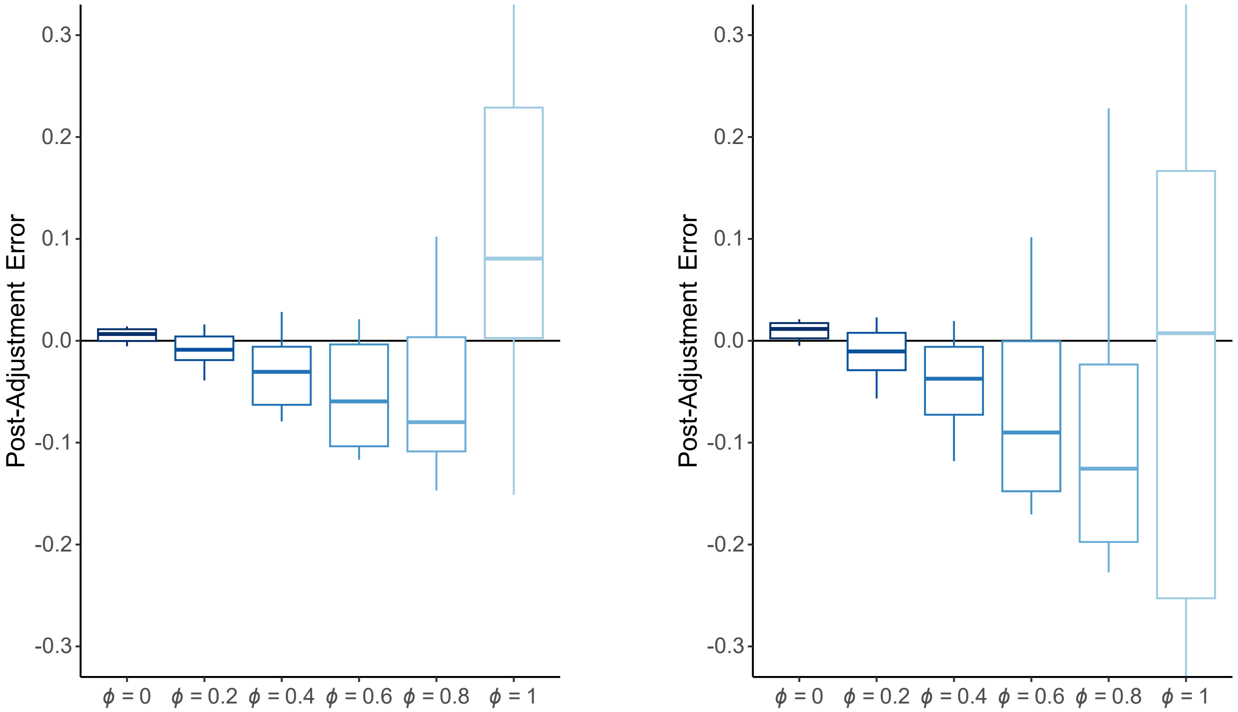


*Note*: Box-plots of error in AM-adjusted genetic correlation estimates across four genetic architectures (full pleiotropy/$r_g$ = 0, full pleiotropy/ $r_g$ = 0.5, partial pleiotropy, and no pleiotropy) and three AM mechanisms (Single-trait AM, Cross-trait AM, and Multi-trait AM). Each architecture/AM-mechanism combination was simulated 20 times. Full details on these simulations are found in the Online Methods. The parameter $\phi$ represents the fraction of the variation in the PGI that is due to error. **(a)** Generation 2 **(b)** Generation 3

**Figure S7: Simulated Effect of AM on Mendelian Randomization**

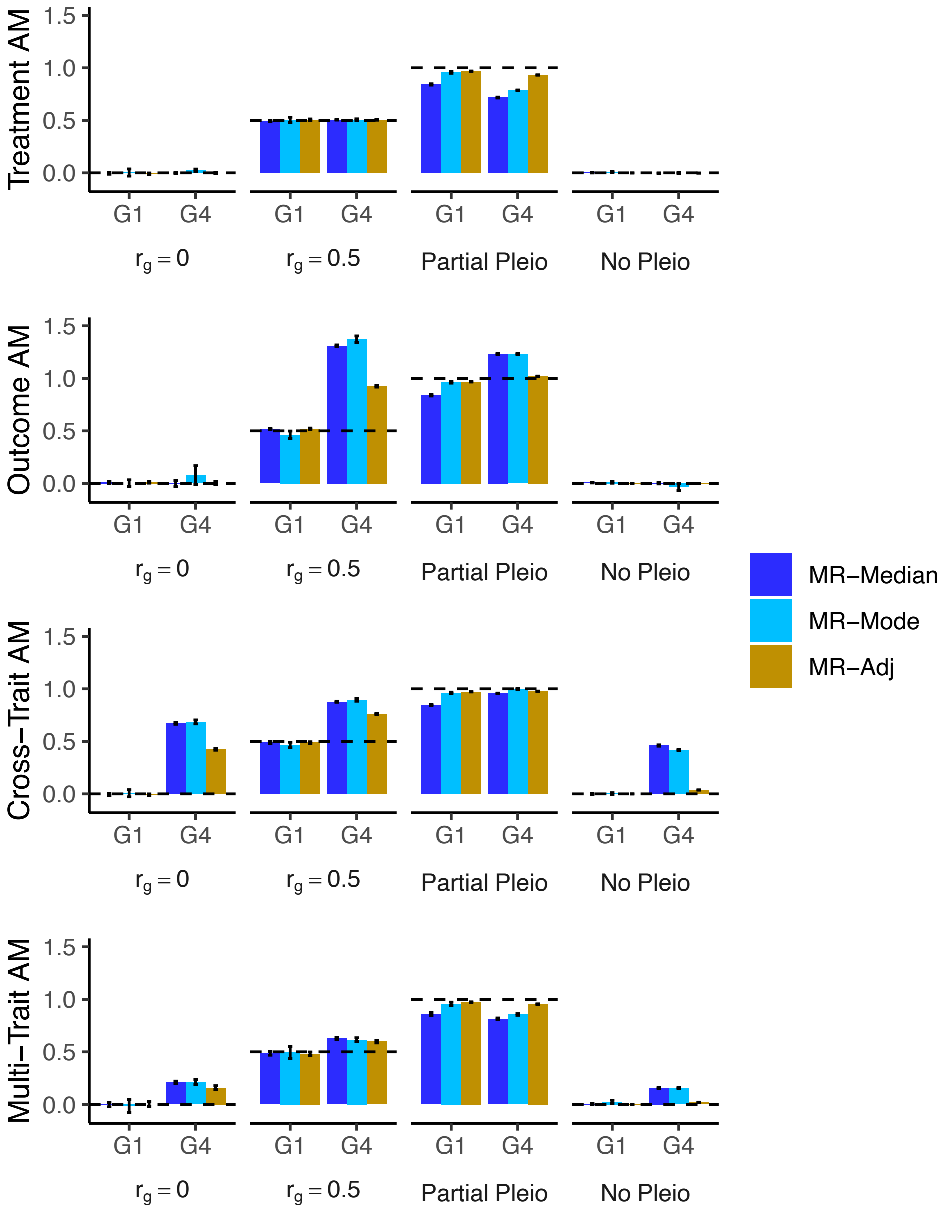


*Note*: This figure shows the simulations results for the effect of AM on various Mendelian Randomization estimators. We consider four different genetic architectures: total pleiotropy/zero correlation (***$r_g$*=0**), total pleiotropy/positive correlation (***$r_g$*=.5**), partial pleiotropy (**Part Pleio**), and no pleiotropy (**No Pleio**). We evaluate the genetic correlation in the first generation (**G1**), which is modeled under a random mating assumption, and in the fourth generation (**G4**), which corresponds to a population undergoing three generations of AM. We model sorting on the treatment treat (**Treatment AM**), sorting on the outcome trait (**Outcome AM**), **Cross-trait AM**, and **Multi-trait AM**. In the multi-trait AM case, people sort on a linear combination of both traits with a weight of 0.8 on the treatment trait and 0.2 on the outcome trait. In each replication of the simulation, we simulate a genome with 2000 SNPs even distributed across 20 chromosomes for 10,000 people. We perform 20 replications. The dark blue bars show the mean **MR-median** estimates, the light blue bars show the mean **MR-mode** estimates, and the gold bars show the AM-adjusted MR-median estimates (**MR-adj**) over the 20 replications. The PGIs used in the adjustment are simulated with no error. The dotted line above each set of bars corresponds to the expected MR estimate in a randomly mating population. The error bars correspond to the 95% confidence interval for the mean.

**Supplementary Figure S8: Summary of Error in AM Adjusted MR Estimates**

**(a)** Generation 2

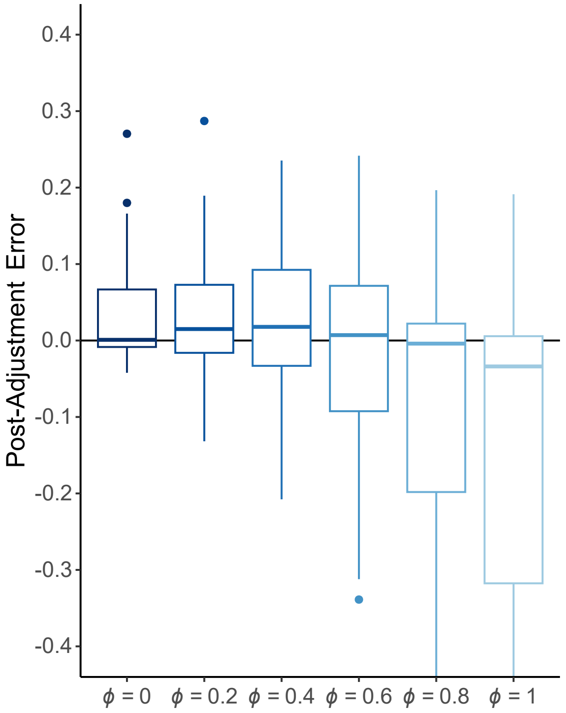


**(b)** Generation 3

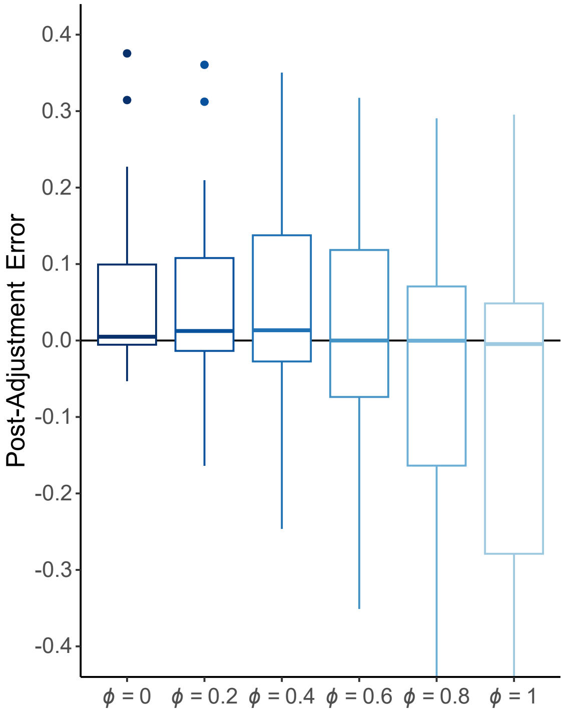


**(c)** Generation 4

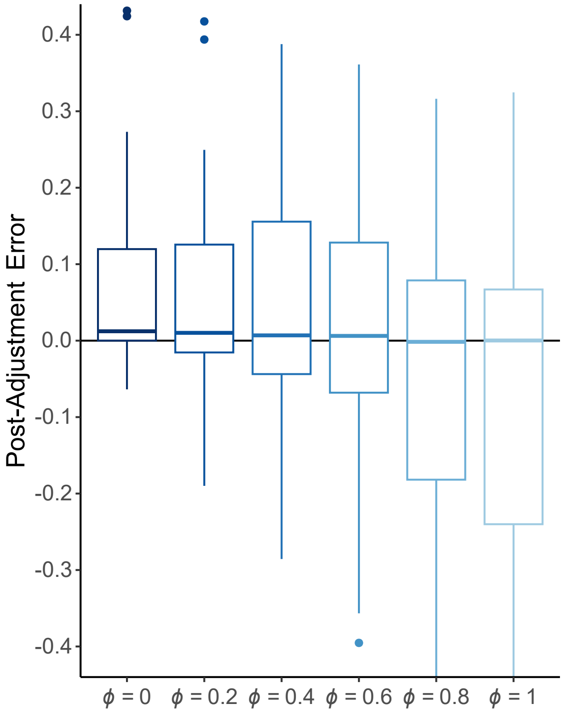


*Note*: Box-plots of error in AM-adjusted median-based MR estimates across four genetic architectures (full pleiotropy/$r_g$ = 0, full pleiotropy/ $r_g$ = 0.5, partial pleiotropy, and no pleiotropy) and four AM mechanisms (Treatment AM, Outcome AM, Cross-trait AM, and Multi-trait AM). Each architecture/AM-mechanism combination was simulated 20 times. Full details on these simulations are found in the

Online Methods. The parameter $\phi$ represents the fraction of the variation in the PGI that is due to error. **(a)** Generation 2 **(b)** Generation 3 **(c)** Generation 4.

**Figure S9: Comparison of Median and AM-adjusted MR Estimates for the Effect of EA on Health**

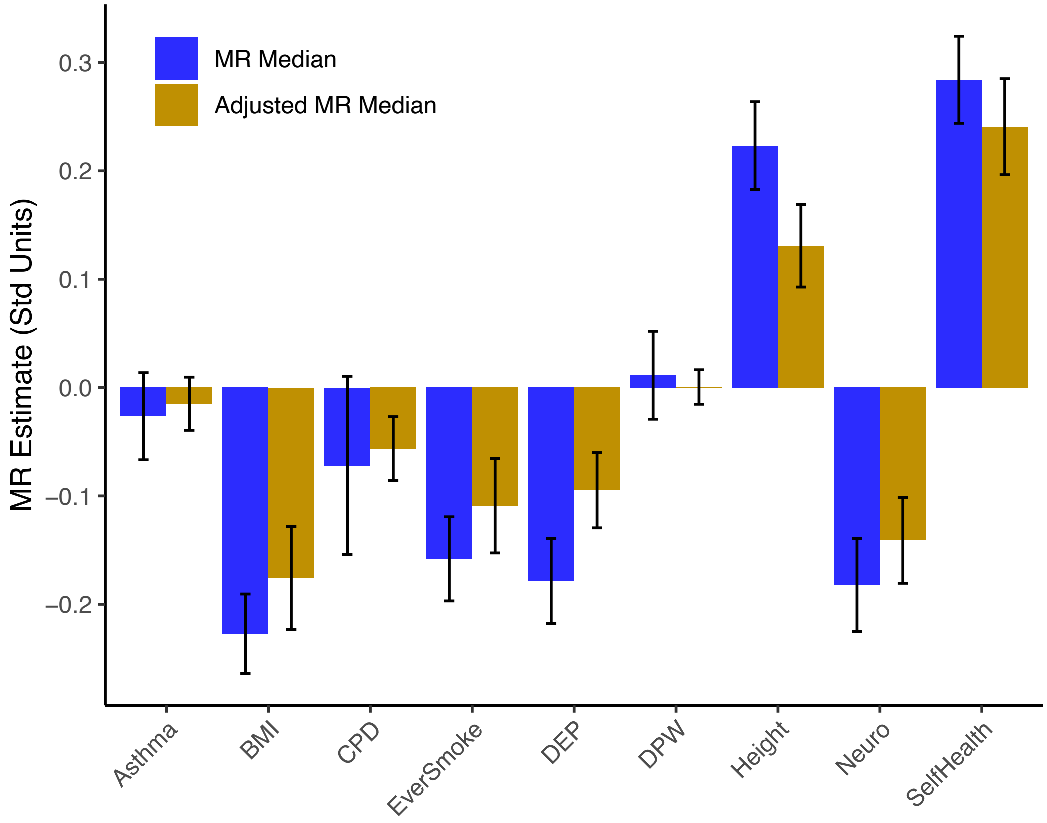


*Note*: Unadjusted (blue bars) and adjusted (gold bars) median-based MR estimates for the estimated causal effect of educational attainment on the specified health trait. Data for this analysis comes from a subset of the UK Biobank (*N* = 148,506) and the PGIs come from the SSGAC's PGI Repository (Becker et al. 2021). BMI: Body mass index; CPD: Cigarettes per day; DEP: Depressive symptoms; DPW: Drinks per week; Neuro: Neuroticism; Self Health: Self-rated health.
